# Phylogenomic reconstruction of *Cryptosporidium* spp. captured directly from clinical samples reveals extensive genetic diversity

**DOI:** 10.1101/2024.04.17.589752

**Authors:** A. Khan, E.V.C. Alves-Ferreira, H. Vogel, S. Botchie, I. Ayi, M.C. Pawlowic, G. Robinson, R.M. Chalmers, H. Lorenzi, M.E. Grigg

**Author notes:** Contributed equally (A. Khan, E.V.C. Alves-Ferreira). Current Address: Animal Parasitic Diseases Laboratory, Agricultural Research Service, USDA, Beltsville, MD 20705, USA.

## Abstract

*Cryptosporidium* is a leading cause of severe diarrhea and mortality in young children and infants in Africa and southern Asia. More than twenty *Cryptosporidium* species infect humans, of which *C. parvum* and *C. hominis* are the major agents causing moderate to severe diarrhea. Relatively few genetic markers are typically applied to genotype and/or diagnose *Cryptosporidium*. Most infections produce limited oocysts making it difficult to perform whole genome sequencing (WGS) directly from stool samples. Hence, there is an immediate need to apply WGS strategies to 1) develop high-resolution genetic markers to genotype these parasites more precisely, 2) to investigate endemic regions and detect the prevalence of different genotypes, and the role of mixed infections in generating genetic diversity, and 3) to investigate zoonotic transmission and evolution. To understand *Cryptosporidium* global population genetic structure, we applied Capture Enrichment Sequencing (CES-Seq) using 74,973 RNA-based 120 nucleotide baits that cover ∼92% of the genome of *C. parvum*. CES-Seq is sensitive and successfully sequenced *Cryptosporidium* genomic DNA diluted up to 0.005% in human stool DNA. It also resolved mixed strain infections and captured new species of *Cryptosporidium* directly from clinical/field samples to promote genome-wide phylogenomic analyses and prospective GWAS studies.

## Introduction

Diarrheal diseases account for about half a million child fatalities every year worldwide, making them one of the leading causes of morbidity and mortality in children less than five years of age (1,2). Although most diarrhea-related deaths are preventable through adequate sanitation, children with impaired immunity, malnutrition, or immunocompromised adults are at the most risk of life-threatening diarrhea. Strikingly, based on a recent Global Enteric Multicenter Study (GEMS) (1), the intestinal apicomplexan parasite *Cryptosporidium* is the second leading cause of severe diarrhea in infants under two years of age and is the most common persistent diarrhea-causing pathogen in young children in Africa and southern Asia (3). In addition to diarrhea-related death, cryptosporidiosis in children is associated with malnutrition, persistent growth retardation, impaired immune responses, and cognitive deficits (4,5). Due to the absence of vaccines, the control of cryptosporidiosis relies solely on sub-optimal chemotherapy, which is limited in efficacy and supply (6). Thus, there is an urgent need for new genetic technologies to identify potential drug/vaccine targets to combat this parasitic infection.

Currently, greater than 20 *Cryptosporidium* species have been identified to infect humans (7,8). Genotyping markers that are applied to resolve the species causing infection include conserved genes such as the 18S small subunit (SSU) rRNA, the 70 kDa heat shock protein (*hsp70*), actin, and the oocyst wall protein 1 (*cowp1*) genes (9). The most commonly used subtyping target is the highly polymorphic 60 kDa glycoprotein gene (*gp60*). The major human cryptosporidiosis-causing pathogens are *C. hominis* and *C. parvum* (10). Despite their high genetic similarity, *C. hominis* maintains a strictly anthroponotic transmission cycle whereas a majority of *C. parvum* genotypes are considered zoonotic pathogens. In addition to these two species, several additional zoonotic species have been detected infecting humans including *C. canis* (usual host, dogs), *C. cuniculus* (rabbits), *C. felis* (cats), and *C. meleagridis* (various) (11). The genetic basis for host adaptation, zoonotic and/or anthroponotic transmission is unclear due principally to a paucity of *Cryptosporidium* spp. whole-genome sequence (WGS) data from this parasite’s wide host range.

Over the last decade, WGS data using next-generation sequencing (NGS) has grown extensively, however it has not been widely applied to *Cryptosporidium* because it is difficult to purify enough parasite material from clinical samples or propogate the parasite *in vitro*. Specifically, few animal models are available to propogate parasites, *and in vitro* propagation is not well developed to produce sufficient pure genomic DNA for WGS (12–14) (15). Additionally, the current oocyst purification efforts to extract high quality gDNA involve sucrose flotation (16), discontinuous sucrose gradient centrifugation (17), cesium chloride (CsCl) gradient centrifugation (18), and immunomagnetic separation (IMS) (19). These steps to enrich parasite oocysts must be performed using fresh clinical/field samples, which must be collected and stored without freezing because the freeze-thaw procedure mechanically ruptures the oocyst wall rendering the techniques above intangible (20). Third, the absence of autofluorescence by *Cryptosporidium* oocysts makes it difficult to FACS-purify single oocysts to perform single-cell sequencing. Finally, current enrichment methods often copurify material with similar buoyant-density or substances adhered to the surface of the oocysts, including host cells (>3Gb) and food particles (>300Mb), that significantly reduce sequence coverage of the *Cryptosporidium* genome (20).

Currently, the few WGS studies pursued to investigate *Cryptosporidium* have been conducted by concentrating and purifying oocysts from stool samples (21–24). However, these are typically limited to symptomatic samples that possess relatively high parasite burdens (≥10^3^ oocysts per gram). Strikingly, asymptomatic clinical manifestation among African and South Asian children is very common (25) and parasite oocyst numbers in these patients are very limited. High-resolution WGS followed by comparative genomics between asymptomatic versus symptomatic carriers is critical to address how parasite genetic factors may influence cryptosporidiosis in the context of other variables such as host immunity and genetics, malnutrition and the gut microbiome. Little progress has been made to develop a highly sensitive WGS technology directly from asymptomatic patient stool samples as most of the enrichment methods result in significant oocyst loss (26). Additionally, it is well documented that increased disease severity has been strongly associated with mixed infections among other closely related apicomplexan parasites such as *Plasmodium* (27). Mixed infection in *Cryptosporidium* has been shown to promote genetic exchange, both intra- and inter-specific in the evolution of new subtypes that possess different biological potentials or host preferences (28). Specifically, admixture of these highly genetically diverse strains has led to the spread of drug-resistant parasites and the emergence of new strains with altered host preferences, or the creation of hypervirulent strains (29–33). High-resolution WGS data has proved to be highly sensitive and discriminatory and can detect population heterogeneity among circulating pathogens (*i.e.* mixed infection) by deconvoluting individual genotypes within a sample (34–36). Thus, it is extremely critical to understand the prevalence of mixed infection using WGS data in high endemic areas and its role introducing genetic diversity through recombination among currently circulating *Cryptosporidium* strains (21).

Capture enrichment sequencing (CES-Seq) has been applied successfully to concentrate *Wolbachia* DNA from whole insect DNA extracts (37), to enrich *Yersinia pestis* DNA from Black Death victims (38), and to detect and characterize felid pathogens for veterinary diagnosis and discovery (39). Additionally, RNA enrichment methods for high resolution quantitative transcriptional analysis using SureSelect CES-Seq have been used *in vivo* to enrich fungal transcripts up to 1,600-fold (40). Recently many genomes of *Leishmania donovani* were sequenced directly from visceral leishmaniasis patient samples and the isolates sequenced *in situ* possessed lower aneuploidy and fewer genomic differences than culture-derived amastigotes from the same patients (41). Our aim was to develop *Cryptosporidium* WGS by applying CES-Seq directly on DNA extracted from clinical stool samples to advance the field of *Cryptosporidium* population genetics, to identify the extent to which mixed infections occur and to identify the true diversity of *Cryptosporidium* species that infect people throughout the world, whether or not symptomatic disease is reported. To perform WGS directly from stool samples, without the requirement to purify oocysts directly from fresh samples, we developed 74,973 RNA baits (probes) to capture, amplify and sequence the whole genome of *Cryptosporidium* using SureSelect technology (Agilent, CA, USA). We demonstrate that the probes can successfully capture as little as 0.005% target gDNA present in a clinical sample at WGS resolution, and that the method captures, with nearly equal efficiency, the medically important species *C. parvum*, *C. hominis,* and *C. meleagridis.* We also show that the CES-Seq method resolved infections with other more distantly related *Cryptosporidium* species, including *C. ubiquitum* and *C. canis*, can be applied to previously frozen material, and is able to distinguish multiple genomes in artificially mixed clinical samples at WGS resolution.

## Materials and Methods

### Ethics Statement and Clinical Samples

Stool samples collected from individuals from Colombia, Ecuador and Egypt were analyzed for the presence of *Cryptosporidium* DNA. These samples had been examined previously for the presence of other protists, as described previously (42). DNA extracted from stool samples from *Cryptosporidium* positive patients from collaborators in Ghana and the United Kingdom were also investigated. Briefly, the cohort population from Colombia consisted of healthy volunteers (n=79) between 16 to 41 years-old from equatorial Colombia who had confirmed *Giardia* infections by microscopy. Fecal samples from these volunteers were preserved in 100% ethanol. The second cohort populations were from Ecuador (n=12) and Egypt (n=24) and were also comprised of microscopically *Giardia* positive patient samples that we assayed for co-infection with *Cryptosporidium* spp. These samples were collected based on approved protocols by the ethics committee of Universidad INCCA de Colombia (protocol number = 237894) with written consent from volunteers and patients, as described previously (42). Ethics approval to obtain clinical samples from Ghana (n=10) was obtained from the Noguchi Memorial Institute for Medical Research Institutional Review Board (Certified Protocol Number: 061/15 - 16). Permission was also obtained from the appropriate authorities from each study site. DNA from clinical samples from the UK (n=10) was provided by Public Health Wales – PHW – Microbiology and Health Protection, UK under a material transfer agreement between NIH and PHW for the analysis of de-identified *Cryptosporidium* DNA, which did not require ethical approval. DNA was extracted from patient samples according to the protocols listed below, anonymized by dis-linking all patient identifiers, and the extracted gDNAs were shipped to NIH for CES-seq.

### DNA Isolation

Genomic DNA was extracted from *C. parvum* oocysts (purchased from Bunchgrass Farms) and the UK clinical samples using the Qiagen DNA Stool kit whereas for all other clinical samples, the Dneasy PowerSoil Pro kit (Qiagen, USA) was used, according to the manufacturer’s instructions (Supplemental figure 1A). Specifically, 1 × 10^7^ excysted oocysts of *C. parvum* purchased from Bunchgrass Farms were utilized to prepare the gDNA which was used to generate artificial mixtures by diluting *C. parvum* gDNA into gDNA from healthy human stool samples*. C. hominis* DNA from isolate TU502 (NR-2520) and *C. meleagridis* DNA from isolate TU1867 (NR-2521) were obtained from BEI Resources (Manassa, VA). The gDNAs were analyzed by 1% agarose gel electrophoresis, stained with ethidium bromide, imaged by Syngene Gel documentation system (GBX-CHEMI-XL1.4. Fisherscientific, USA) and compared with GeneRuler 1kb Plus DNA Ladder (Thermo Scientific, USA). Quantity and quality of gDNA were assessed using the Invitrogen Qubit 4 Fluorometer (Invitrogen, USA), DS-11 Series Spectrophotometer (DeNoVix, USA), 4200 TapeStation System (Agilent, USA), and 2100 Bioanalyzer Instrument (Agilent, USA).

### Real-time Quantitative PCR (qPCR)

Real-time qPCR was performed using 18S rRNA gene primers (Table S1), a QuantStudio 6 Flex Real-Time PCR system (Applied Biosystems, USA), with a 20 µl reaction mixture, containing 1 X Power SYBR Green PCR master mix (Applied biosystems, USA), 500nM of each primer, and 10-fold serial dilutions (from 10ng to 0.0001ng) of *C. parvum* gDNA. Sterile water and gDNA from healthy volunteers were used as non-template controls and were run with every assay. All assays were performed in triplicate to ensure reproducibility.

### Sanger Sequencing of 18S rRNA Gene for Species Assignment

DNA extracted from clinical samples infected with *Cryptosporidium* were also subjected to 18S rRNA gene Sanger sequencing to assign species after PCR amplification using nested primers: external primers, 18SrRNA(ext)F (5’-CCTGCCAGTAGTCATATGCTTG-3’) and 18SrRNA(ext)R (5’-GAATGATCCTTCCGCAGGT-3’); internal primers, 18SrRNA(int)F (5’-GTTAAACTGCGAATGGCTCA-3’) and 18SrRNA(int)R (5’-CGAAACTTTCCTTACATGTATTGCT-3’). PCR products were purified with a QIAquick PCR purification kit (QIAGEN, USA) according to the manufacturer’s instructions. Sequencing was conducted with two independent templates using BigDye cycle sequencing (Applied Biosystems, USA) by Quintara Bio (Quintarabio, USA). Nucleotide sequences were assembled using FinchTV 1.4 Sequence Alignment Software (https://digitalworldbiology.com/FinchTV, Geospiza, Inc) and aligned with previously published sequences in GenBank (https://www.ncbi.nlm.nih.gov/genbank/) using ClustalX (43) with default settings. Maximum-likelihood phylogenetic trees were generated with the aligned sequences in Nexus format (44) using Molecular Evolutionary Genetic Analysis (MEGA-X) (45), Tamura-Nei model (46). 1000 replicates were used to generate the bootstrap values.

### Whole Genome CES-Seq

Initially, artificial mixtures (*C. parvum* genomic gDNA diluted in gDNA extracted from healthy human stool samples) of 200 ng total gDNA in different ratios were generated for capture-hybridized sequencing using genome-wide biotinylated RNA baits for *Cryptosporidium*. In total, 74,973 RNA baits (probes) were generated based on the available reference genome of *C. parvum* at the time this study was initiated (https://cryptodb.org/common/downloads/release-34/CparvumIowaII/fasta/data/). Duplicated baits and baits with high similarity to the human genome were filtered out. Exclusion criteria for bait selection were known regions of mis-assembly, long stretches of low complexity, and gene-poor regions in telomeric and sub-telomeric regions. In total, baits were designed to capture approximately 92% of the *C. parvum* genome, based on the assembly of the Iowa II isolate, released on CryptoDB_v34 (Table 1). Within the genome regions that were included for bait design, every nucleotide position was covered with at least 2 overlapping baits, except in highly variable regions, including telomeric and sub-telomeric regions that were gene-rich, in which 5 baits were designed.

**Table 1.** Design of *Cryptosporidium* specific biotinylated baits for SureSelect target enrichment sequencing.

| Chromosome<sup>a</sup> | Target Size<sup>b</sup> | Covered<sup>c</sup> | Fraction Covered |
| --- | --- | --- | --- |
| CM000429 | 875659 | 808459 | 0.9232578 |
| CM000430 | 985969 | 902040 | 0.91487663 |
| CM000431 | 1099352 | 1019400 | 0.92727352 |
| CM000432 | 1104417 | 977040 | 0.88466585 |
| CM000433 | 1080900 | 945720 | 0.87493755 |
| CM000434 | 1332857 | 1204320 | 0.9035628 |
| CM000435 | 1278458 | 1210560 | 0.94689071 |
| CM000436 | 1344712 | 1206000 | 0.89684631 |

<sup>a</sup> Genome sequence of *C. parvum* was utilized to develop biotinylated baits throughout the genome (https://cryptodb.org/common/downloads/release-34/Cparvumlowall/fasta/data/CryptoDB-34_Cparvumlowall_Genome.fasta)
<sup>b</sup> Total base pair (bp) size of each chromosome of *Cryptosporidium parvum*.
<sup>c</sup> Total base pair (bp) size of the covered genome using biotinylated baits for SureSelect TES.

Genomic DNA of artificial samples and clinical samples were first sheared by focused ultrasonication (Covaris Inc., USA) into 200 to 300 bp fragments followed by a quality control assessment using the Invitrogen Qubit 4 Fluorometer (Invitrogen, USA), 4200 TapeStation System (Agilent, USA), and 2100 Bioanalyzer Instrument (Agilent, USA). Capture enrichment libraries were prepared using SureSelect^XT^ ^HS^ Target Enrichment System for Illumina Paired-End Multiplexed Sequencing Library (https://www.agilent.com/cs/library/usermanuals/public/G9702-90000.pdf, Agilent Technologies, USA) using genome wide baits with 12 amplification cycles following the manufactures’ protocol (Fig. 1A). Amplified capture libraries were purified with streptavidin coated beads and quantified again with Bioanalyzer DNA 1000 chip (Agilent, USA) and D1000 ScreenTape (Agilent, USA). Finally, the genomic libraries were prepared with Nextera XT Kits (Illumina, USA) and DNA sequencing was performed using paired-end reads on a MiSeq, NextSeq or HiSeq system (Illumina, USA).

**Fig. 1.**
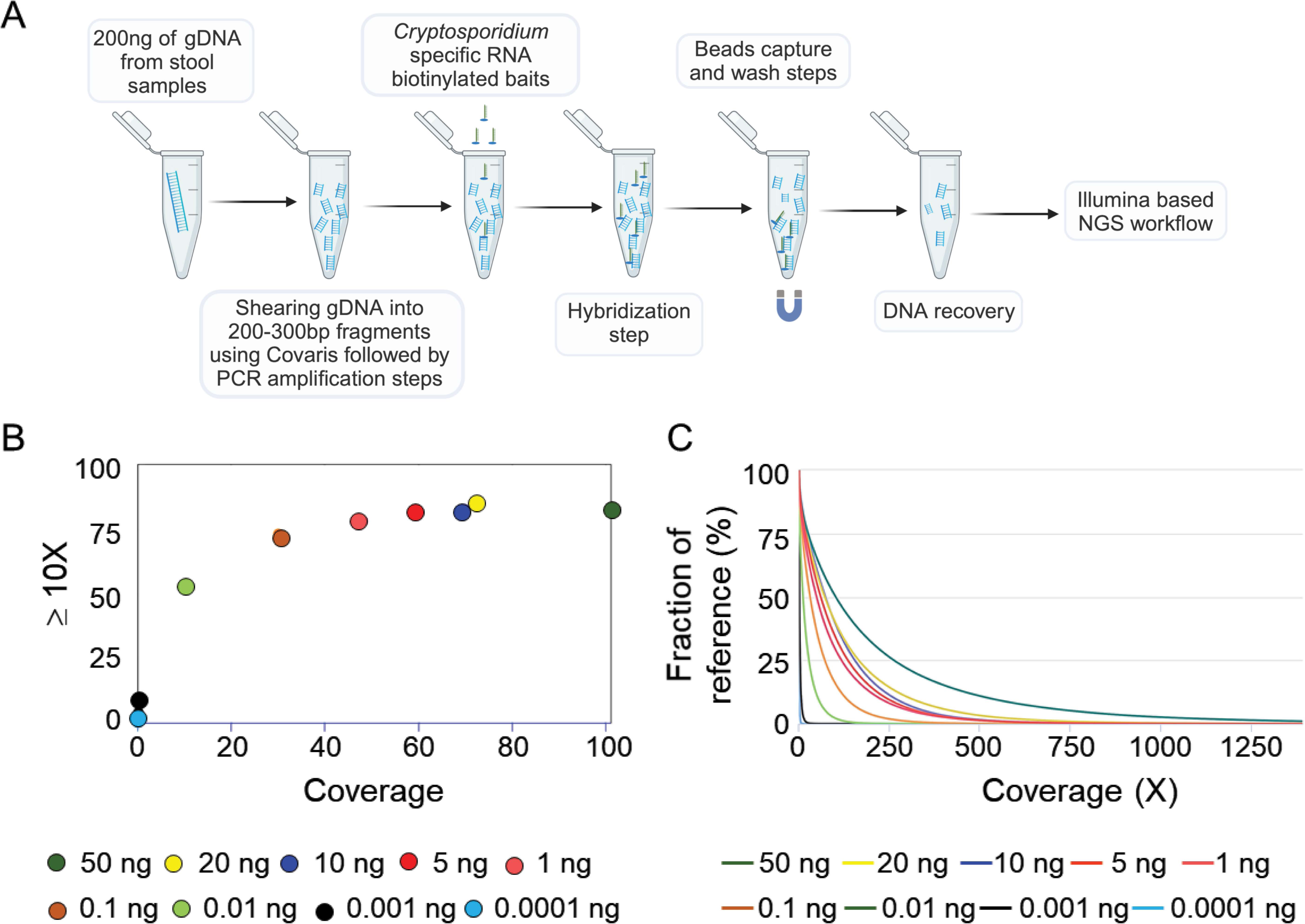
Validation of SureSelect CES of protozoan diarrheal agent *Cryptosporidium.* A) Workflow of SureSelect TES. Whole genome of *Cryptosporidium* was captured by hybridization with genome wide uniformly distributed biotinylated probes (Table 1). The strand extension and the capture steps allow the recovery of the amplified library for NGS using Illumina sequencing. B) Graphical display of average sequencing depth and coverage percentage with ≥ 10X for SureSelect capture enrichment followed by whole genome sequencing using Illumina platform. Different dilutions of *C. parvum* gDNA in 200 ng of total gDNA are plotted with different color circles as listed in the legend. C) Fraction of the reference genome recovered after target enrichment using genome wide RNA baits and whole genome sequencing using Illumina platform. Different dilution of *C. parvum* gDNA are depicted with different color lines as listed in the legend.

### DNA Sequence Read Processing

Paired-end reads from CES-Seq were trimmed using Trimmomatic v.0.36 (47) to remove sequencing adapters and low quality bases, followed by filtering of reads derived from human contamination. Filtered raw reads were aligned against the *C. parvum* reference genome CryptoDB-54_CparvumIowa-ATCC_Genome (Table S3) using the Burrows-Wheeler Aligner (BWA mem, v0.7.15) (48). BAM files were processed with SAMtools v1.3.1(49), BEDtools v2.25.0 (50), and Picard tools v2.7.1 (http://broadinstitute.github.io/picard) to remove duplicated reads. Reads were then locally realigned around potential INDELs with GATK v3.7 tools RealignerTargetCreator and IndelRealigner. Quality control metrics of the whole genome sequenced data were quantified and visualized by MultiQC (51). Next, SNP data was generated from processed bam files using a customized SNP annotation pipeline. Briefly, raw SNPs and INDELs were identified with HaplotypeCaller followed by GenotypeGVCFs tools from the GATK (52). All SNPs having either QD (Quality of Depth) < 2, FS > 60, MQ <40, MQRRankSum < −12.5, or ReadPosRankSum < −8.0 were filtered out. VCF files containing the SNPs that passed the filtering step were further filtered to select only those SNPs that were supported by a minimum of 5 reads in at least one of the samples present in each VCF file.

### Genetic Diversity and Structural Variation Analyses

Filtered genome-wide SNPs that differ from the reference (see Table S3) were identified using VCFtools (53), and were concatenated and converted into a table file containing predicted genotypes (one column per sample, one row per SNP position) using GATK tool VariantsToTable (v4.2.0.0). The table was then converted into different multiFASTA files containing pseudosequences from concatenated SNPs for a selected subset of samples with the shell script run_tbl2fasta_loop.sh, which converts the table to FASTA format in order to generate a phylogenetic neighbor-net network tree using SplitsTree v.4.17.1 (54) with 1,000 bootstrap replicates. The total number of SNPs and the nucleotide diversity (ρε) (55) per 10 kb windows were also calculated using VCFtools and were plotted using the circos software (http://circos.ca/) (56). To calculate copy number variation (CNV), the coverage at each base pair was first calculated using genomeCoverageBed, BEDTools (50) and then combined into 10 kb sliding windows using custom scripts as described previously (57). Mean and standard deviations were calculated for each strain and bins were generated as “1X” if the value was up to 1 standard deviation from the mean (1SD) and plotted using ggplot2 package (58) as described previously (57). The allele composition in each variant SNPs per strains was plotted using a bottle brush plot as described previously (59). For the SNP density statistic plots, read depth statistics were calculated with SAMtools depth tool version 1.9 with the-aa parameter on. The remaining SNP statistics were estimated using an in-house python script to present the median SNP density per chromosome, read depth and coverage. Read coverage per 10 kb was estimated as the number of bases in a 10kb window having a read depth of 1X or higher. To account for the presence of sequencing gaps, normalized SNP density was defined as the number of SNPs counted in a 10kb window multiplied by 10,000 and divided by the read coverage of the same window. Dots were used to represent normalized SNP density values (number of high confidence SNPs / 10kb), Green lines to depict read depth / 10kb, Gray lines to represent read coverage / 10kb, and red dashed lines to show median SNP density per chromosome.

### DEploid to Infer Mixed Infections

To create an artificially mixed sample, 1 ng of each *C. parvum*, *C. hominis*, and *C. meleagridis* gDNAs were added to 197 ng of gDNA of human stool sample (total = 200 ng) followed by whole genome sequencing using CES-Seq methodology as described above. The number of strains and their relative proportion and the haplotypes present in the mixed sample were estimated by deconvolving multiple genome sequences using the software package DEploid (35). We downloaded 12 *C. parvum*, 8 *C. hominis*, 2 *C. meleagridis*, 2 *C. ubiquitum,* and 1 *C. tyzzeri* genomes from SRA (https://www.ncbi.nlm.nih.gov/sra) (Table S4) followed by variant calling using GATK (52) to construct the reference genome. SNP positions containing heterozygote genotypes or missing data were removed from the PANEL file, which was used for recalibrating the VCF file of artificial mixed infection samples. DEploid was run in IBD (identity by descent) mode and the relative composition of heterozygous and homozygous SNPs present in the artificial mixed sample was calculated in 10kb sliding windows using custom Java scripts (59) to construct histogram plots in circos (56).

### Population Genetic Structure and Admixture Analyses

Population genetic structure and admixture analyses were determined using the POPSICLE pipeline (60) with 10 kb window block sizes and using the number of clusters set from K=1 to 15 followed by calculating the optimal number of ancestries using the Dunn index (61) as described previously (60). The contribution of each haplotype was then painted in a circos plot (56) with color assignment based on the number of clusters (ancestries). For haplotype plots, we used an in-house python script to calculate the percentage of different SNPs within a non-overlapping sliding window of 10kb between the strain of interest (*i.e.* Isolate EC1, UKUB17 or NMIMR11) and the reference strains of either *C. parvum*, *C. meleagridis*, *C. hominis* or *C. ubiquitum*. Within each of these windows, SNPs falling within sequencing gaps were ignored in the calculation.

## Results

### Cryptosporidium Capture Enrichment (CES-Seq) for Whole-Genome Sequencing

To capture and sequence *Cryptosporidium* at whole-genome resolution directly from stool samples, 74,973 CES-Seq RNA baits were developed, tiled end-to-end to cover ∼92% of the 8.2 Mb *C. parvum* reference genome CryptoDB-34_CparvumIowaII_Genome. The baits were tiled without gaps and with two overlapping sequences, except within hypervariable regions, particularly telomeric and sub-telomeric regions, where at least 5 overlapping sequence baits were used. Regions that were comprised primarily of low complexity and repetitive regions were removed. Genome coverage of the final bait set is listed in Table 1, each chromosome had a different coverage and this ranged from as low as 87.5% to as high as 94.7% (Table 1). To test the sensitivity and specificity of the CES-Seq method, different concentrations of *C. parvum* gDNA (50ng to 0.0001ng) were spiked into human stool gDNA to generate a 200ng artificial gDNA mixture (Fig.1). DNA extraction using the DNeasy PowerSoil Pro kit was determined empirically to produce high yield, high molecular weight DNA preps to optimize shearing of gDNA into 200-300 bp fragments using the Covaris platform (Suppl. Fig. 1). As a guide to identify viable clinical samples suitable for the CES-Seq method, a qPCR assay at the 18S rRNA gene was developed and applied against the artificial mixed gDNA samples to determine the threshold C_T_ associated with high-quality whole-genome sequencing. *Cryptosporidium parvum* gDNA was serially diluted tenfold in human stool gDNA from 10 ng/µl to 0.0001 ng/µl, followed by qPCR (Suppl. Fig 1B and C).

We first determined the lower limit for optimal genome wide analyses using the CES-Seq method against the artificial mixtures of *C. parvum* DNA, as well as capture enrichment efficiency, the percentage of reads that aligned, and the depth of coverage after trimming and quality filtering paired-end reads that aligned against the *C. parvum* Iowa II reference genome. The percentage of aligned reads was greater than 98% and depth of coverage >200X for all artificial mixtures that contained 5ng or greater of *C. parvum* gDNA, presumably due to saturation of the baits at high concentrations of target gDNA (Table 2). Notably, 0.01ng of *C. parvum* gDNA (0.005%), which corresponded to a C_T_ of 24.36, was optimal for genome wide analysis with at least 38X coverage of the reference genome, at the depth of sequencing used (Table 2). Further, linearity assays based on titrating DNA showed an ∼1,500-fold enrichment from a starting concentration of 0.1 ng of *C. parvum* gDNA spiked into 200 ng of total human stool gDNA. Hence, C_T_ values equal to or lower than 24 (0.01 ng/µl *C. parvum* gDNA) were determined empirically as optimal for achieving greater than 30X genome coverage for SNP calling and genotyping at WGS resolution (Fig.1B, C) (Table 2). However, *C. parvum*-specific reads were also recovered at 0.001ng of input DNA (C_T_ of 28), and represented ∼28 Mb of *C. parvum* DNA sequenced that was distributed genome-wide. Although it was considered lower-confidence because the majority of sequences obtained were at only 2-3X coverage, it nevertheless represented a significant improvement over MLST-based genotyping strategies that recover sequences for only 5-10kb of the genome.

**Table 2.** Whole genome sequencing of libraries after amplification using SureSelect Target enrichment sequencing.

| Amount of *C. parvum* gDNA<sup>a</sup> | Total number of sequenced reads<sup>b</sup> | Total number of mapped reads<sup>c</sup> | % of aligned read | Coverage (X) | Insert size | qPCR C<sub>T</sub> |
| --- | --- | --- | --- | --- | --- | --- |
| 5.0ng | 9224038 | 9018315 | 97.8 | 213 | 167 | n.d. |
| 1.0ng | 8774746 | 8097021 | 92.3 | 172 | 166 | 17.35 |
| 0.1ng | 6308564 | 4794797 | 76 | 108 | 156 | 20.80 |
| 0.01ng | 6836298 | 2100385 | 30.7 | 38 | 153 | 24.36 |
| 0.001ng | 5564782 | 201378 | 3.6 | 2 | 138 | 27.99 |
| 0.0001ng | 2256618 | 27146 | 1.2 | ~0.0 | 20 | 30.99 |

<sup>a</sup> Amount of *C. parvum* gDNA spiked into 200ng of total gDNA from host stool samples
<sup>b</sup> total number of sequenced reads after Illumina sequencing using NextSeq500/550 Mid Output Kit v2.5 (300 cycles)
<sup>c</sup> Total number of mapped reads after ref-mapping Illumina short read sequences against reference *C. parvum* reference sequence https://cryptodb.org/common/downloads/release-34/Cparvumlowall/fasta/data/CryptoDB-34_Cparvumlowall_Genome.fasta n.d = not done

### Suitability of *C. parvum* Probes to Capture Other *Cryptosporidium* Species Genomes

To test the efficiency of the *C. parvum* baits to capture other medically important *Cryptosporidium* species at genome-wide resolution, we generated artificial gDNA samples by adding 1 ng of either *C. parvum*, *C. hominis* or *C. meleagridis* gDNA into 199 ng of human stool gDNA. We next submitted these different *Cryptosporidium* species gDNA samples to the capture hybridization protocol to produce libraries that were sequenced using the Illumina platform (Fig. 2A). All paired-end reads were filtered and aligned against the *C. parvum* reference genome for variant calling using Genome Analysis Toolkit (GATK) (52). 37,367 total single nucleotide polymorphisms (SNPs) were identified between *C. parvum* and *C. hominis,* and 345,437 between *C. parvum* and *C. meleagridis*. The SNPs were distributed evenly throughout the genome (Fig. 2B) and were used subsequently for phylogenomic analysis (Fig. 2C). For the phylogenomic comparison, we downloaded two previously published WGSs for each species from SRA database (*C. hominis*: ERR311205, ERR311209; *C. parvum*: ERR1738341, ERR1035621; and *C. meleagridis*: SRR1179185, SRR793561). Additionally, *C. cuniculus* (SRR6813608) and *C. tyzzeri* (SRR5683558) WGSs were analyzed to generate an unrooted phylogenetic tree. CES samples clustered together with publicly available WGSs, *C. parvum*, *C. hominis* or *C. meleagridis* gDNA, establishing the feasibility of applying the *C. parvum*-derived baits against more distantly related *Cryptosporidium* parasites to genotype samples at WGS resolution, such as *C. meleagridis* (Fig. 2C).

**Fig. 2.**
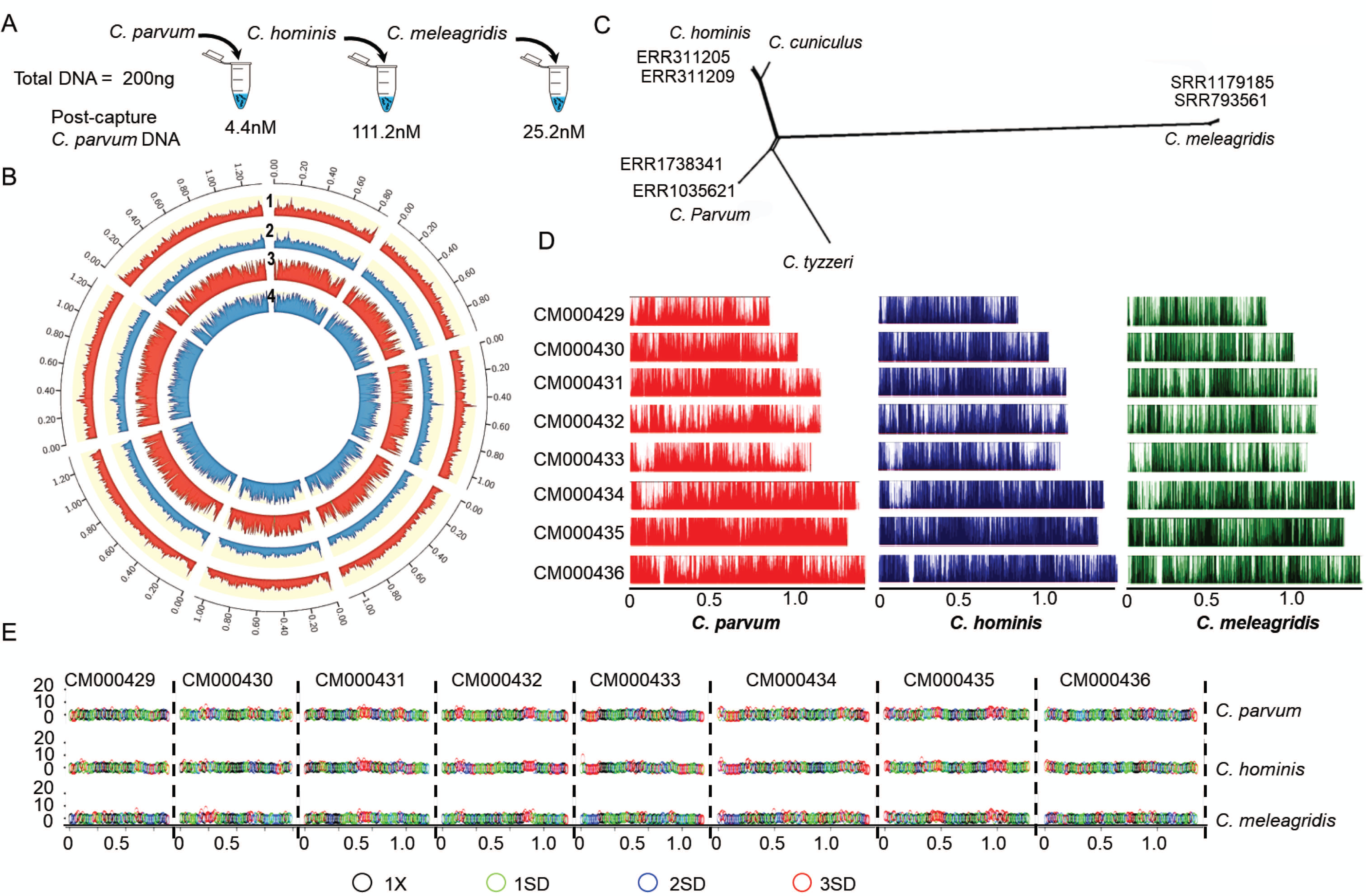
Capture enrichment sequencing of distantly related species of *Cryptosporidium*. A) Diagram to depict SureSelect sequencing of *C. parvum*, *C. hominis*, and *C. meleagridis*. 1 ng of parasite gDNA was introduced into 199 ng of gDNA from healthy human stool sample (Total = 200 ng gDNA). B) Circos plot showed the distribution of pairwise SNPs (red) and Pi (blue) across the genome per 10kb sliding windows. 1 and 2, pairwise SNP and Pi plots between *C. parvum* and *C. hominis* respectively. 3 and 4, pairwise SNP and Pi plots between *C. parvum* and *C. meleagridis* respectively. C) Neighbour-net analysis of *Cryptosporidium* spp. based on genome wide SNPs after amplifying the whole genome using SureSelect baits and Illumina sequencing followed by reference mapping the genome sequences. *C. parvum*, *C. hominis*, and *C. meleagridis* were whole genome sequenced using CES methodology whereas ERR311205, ERR311209, ERR1738341, ERR1035621, SRR1179185, 793561, *C. cuniculus* (SRR7895182), and *C. tyzzeri* (SRR5683558) were downloaded from NCBI SRA (https://www.ncbi.nlm.nih.gov/sra) site. Scale bar represents the number of SNPs per site. D) Bottlebrush representation of genome wide distribution of amplified reads using CES. X-axis = size of chromosome, y-axis = inferred read depth, normalized across the entire genome. Red = *C. parvum*, Purple = *C. hominis*, green = *C. meleagridis*. E) Genome-wide CNV was determined in 10kb tiling windows. The 10kb blocks with no CNV are plotted as black circles (1X). Green, blue, and red dots indicate 1, 2, and 3 SDs from the mean (1X), respectively. *Y*-axis indicates the CNV

To represent the genome-wide distribution of amplified reads by the CES-Seq method, we generated Bottlebrush plots (59) to display the allele count distribution at each SNP position of *C. parvum*, *C. hominis*, and *C. meleagridis*. The plots showed that the reads had a uniform distribution along all eight chromosomes of these three species (Fig. 2D), closely resembling the SNP distribution plot (Fig. 2B) and confirming the high degree of specificity of CES technology. Next, to evaluate whether the CES-Seq method was able to resolve genome-wide structural variation, copy number variation (CNV) was calculated by estimating the combined coverage/base pair in a 10 kb sliding window. CNV estimation using raw reads from CES samples showed significant changes across the genome within *C. parvum*, *C. hominis* and *C. meleagridis,* particularly in subtelomeric regions (Fig. 2E).

It is often the case, that in endemic regions, individuals with cryptosporidiosis are infected with multiple strains of *Cryptosporidium* that possess varying levels of relatedness (62). To determine the capability of the CES-Seq method to resolve a mixed-species infection, and to detect the relative proportion of the different species present in a sample, we next generated a mock community by adding 1 ng each of *C. parvum*, *C. hominis*, and *C. meleagridis* gDNA to 197 ng of human stool gDNA to perform WGS-CES protocol (Fig. 3A). In order to deconvolute the mixed infection, aligned raw reads were analyzed using DEploid, which uses haplotype structure with a reference PANEL of cloned isolates of defined haplotypes to reference map against in order to calculate allele frequencies (35,36). In this case, the reference PANEL used was comprised of 12 *C. parvum*, 8 *C. hominis*, 2 *C. meleagridis,* 2 *C. ubiquitum*, and 1 *C. tyzzeri* genomes (Table S4). DEploid detected the presence of all three species in the mock mixed sample with a ratio of 43% *C. parvum*, 50% *C. hominis*, and 7% *C. meleagridis* (Fig. 3B). To investigate whether the lower relative proportion of raw reads captured from *C. meleagridis* (at 7%) was related to the higher degree of genetic polymorphism between *C. meleagridis* and *C parvum*, we assessed the genome-wide distribution of heterozygous versus homozygous SNPs in blocks of 5 kb across each chromosome. The hetero-homozygosity sliding window plot (Fig. 3C) corresponded closely with the DEploid estimation, because a significant proportion of SNPs (resolved as blocks of yellow within the circos plot) within the *C. meleagridis* genome plot were filtered out prior to conducting the DEploid analysis. However, the analysis pipeline did establish that *C. meleagridis* specific SNPs were detected with sufficient frequency, with genome-wide distribution, to validate this approach as a viable method to resolve mixed species infections in a biological sample.

**Fig. 3.**
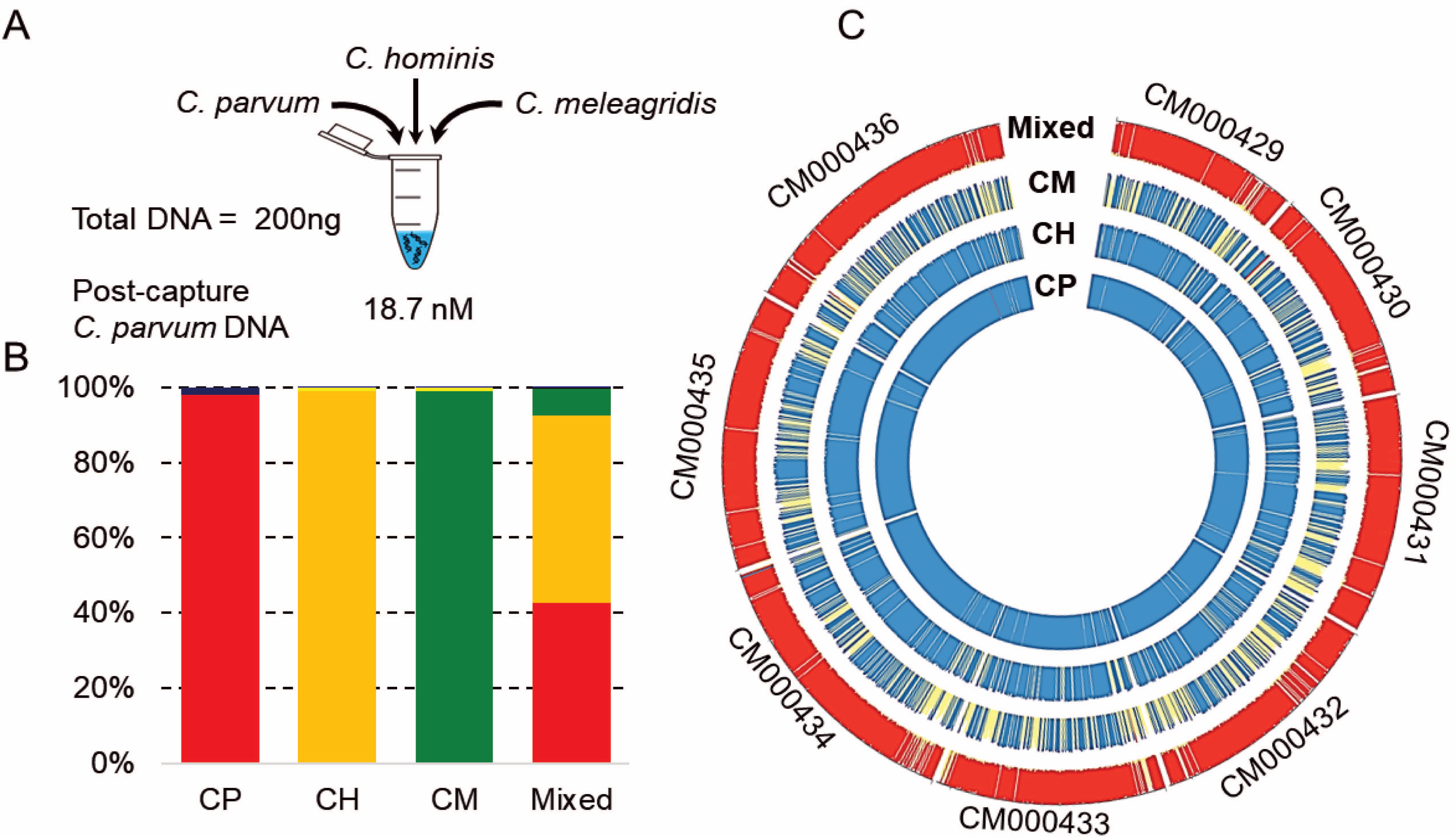
Detection of mixed infection using capture enrichment sequencing. A) Diagram to generate an artificial mixed sample. 1ng of *C. parvum*, *C. hominis*, and *C. meleagridis* gDNAs (in total 3ng of *Cryptosporidium* gDNA) were mixed with 197ng of fecal stool sample (total = 200ng) gDNA to create a mixed infection of sample with the presence of three *Cryptosporidium* species B) Estimation of multiplicity of infection by genome wide abundance of heterozygous genotypes using software Deploid (35). CP = *C. parvum*, CH = *C. hominis*, CM = *C. meleagridis*, Mixed = mock mixed infection sample generated as described previously (Fig 3. A). *X*-axis indicates the percentage of abundance of species in each sample. CP, CH, and CM data were generated based on Fig. 2A. C) Circos representation of genome wide heterozygosity and homozygosity plots of 3 *Cryptosporidium* species and the mixed sample. Red color = >90% of heterozygous SNPs, blue = > 90% of homozygous SNPs, yellow = 50% heterozygous, 50% homozygous SNPs. Each track represents each sample. Chromosome numbers are shown on the outer ring. CP = *C. parvum*, CH = *C. hominis*, CM = *C. meleagridis*, Mixed = mock mixed sample

### Utility of CES-Seq to Assemble *Cryptosporidium* at WGS Resolution Directly from Human Stool Samples

To investigate the capacity of CES-Seq methodology to capture genome-wide *Cryptosporidium* spp. SNP variation directly from previously frozen or ethanol-preserved clinical stool samples without any prior oocyst purification or enrichment among isolates, we tested human stool samples from Colombia (N=79), Ecuador (N=12), and Egypt (N=24) that had been determined previously to be *Giardia*-positive and were obtained from healthy adults with no gastro-intestinal clinical symptoms (42) (Fig. 4A). To determine which of the stool samples were qPCR positive for *Cryptosporidium*, we applied modified GEMS primers (63,64) to screen for the presence of the following three enteroparasites: *Cryptosporidium, Giardia* and *Entamoeba histolytica* (Table S1). All samples except one (ID-2) were positive for *Giardia*, as expected. The presence of *Giardia* gDNA served as an important control to investigate the specificity of the CES-Seq method to show that it specifically pulls out *Cryptosporidium* DNA in the context of a mixed infection with other enteroparasites unrelated to *Cryptosporidium.* Four samples were positive for the presence *of E. histolytica*, 3 from Colombia, and 1 from Egypt, with a prevalence rate of 3.5% (4/115) (Table S2). Four samples gave positive C_T_ values for *Cryptosporidium*, 1 from Colombia, 2 from Ecuador, and 1 from Egypt, with a prevalence rate of 3.5% (4/115) (Table S2). The Colombia sample was only weakly positive for *Cryptosporidium* (COL3, C_T_=39.35) and *Giardia* (C_T_=36.15), but was strongly positive for *E. histolytica* (COL3, C_T_=18.19). Due to the high C_T_ for *Cryptosporidium,* we excluded COL3 but selected EC1, EC4 and FEgypt samples for CES-Seq. Additionally, we screened 20 *Cryptosporidium*-positive samples, 10 from Ghana and 10 from the UK, that had been previously tested by qPCR (Table S2). We selected three human fecal samples from each country, Dg045 (C2), Dg083(C8), NMIMR11, UKP196, UKH101 and UKUB17 for CES-Seq based on their low C_T_ values, the *Cryptosporidium* species expected to be present, and gDNA quality (assessed by TapeStation).

**Fig. 4.**
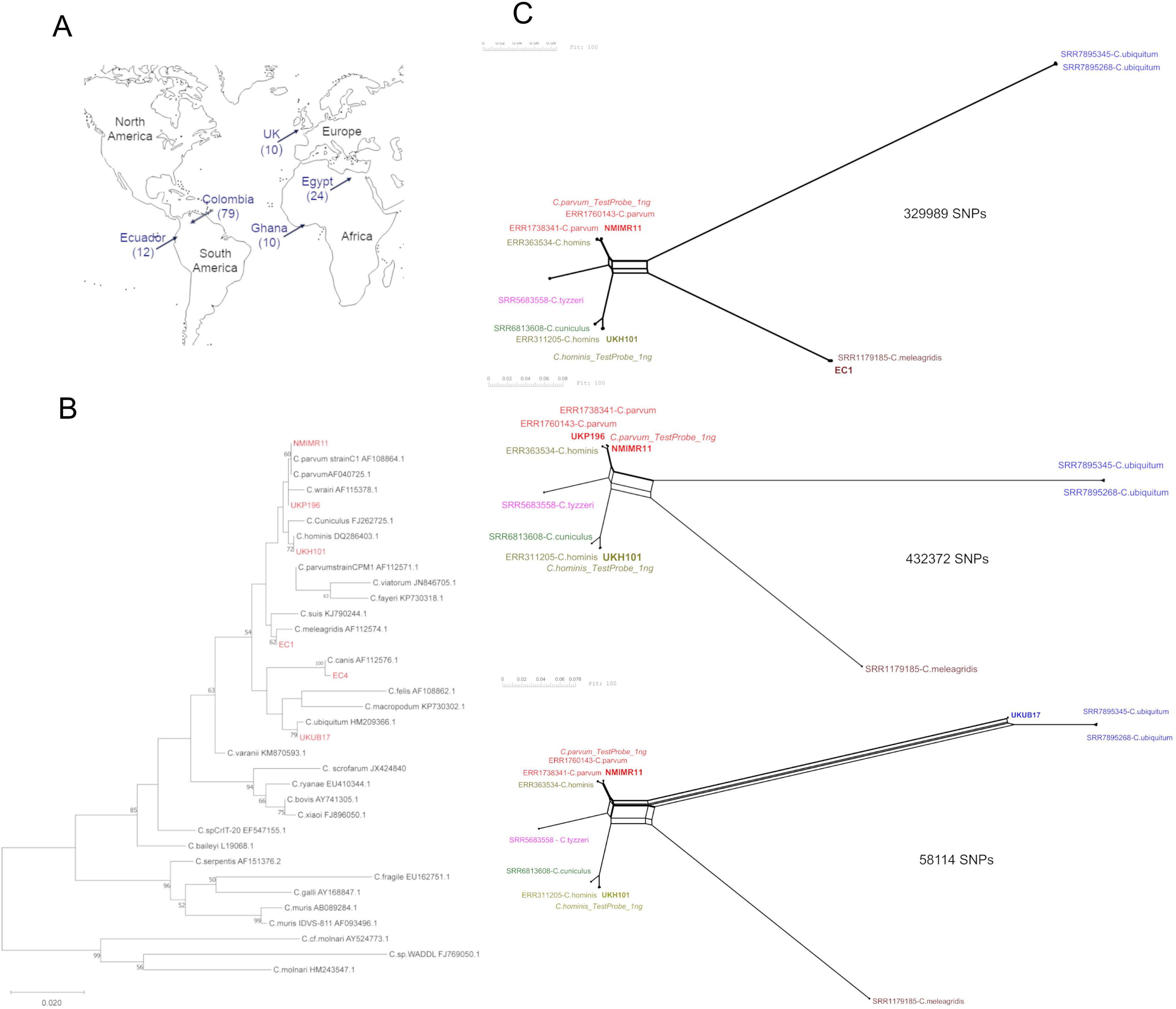
Detection of *Cryptosporidium* in asymptomatic and symptomatic patient’s stool samples. A) Geographic locations of clinical samples used in this study show in arrow heads. Sample numbers are in the parentheses. B) Phylogenetic analysis of 18S rRNA. An unrooted maximum-likelihood tree was generated using Geneious-PhyML tree with 1,000 bootstrap replicates. Representative sequences of different species of *Cryptosporidium* were downloaded from NCBI (https://www.ncbi.nlm.nih.gov/) and compare with the sequenced samples (green). Bootstrap values between 75 to 100% are represented by the red nodes, values between 50 to 75% are depicted by blue nodes, and values between 25 to 50% are denoted by yellow nodes. Scale = number of SNPs per site. C) Phylogenetic Network Trees. Trees were generated with SplitsTree (v4.17.1), total number of SNPs below each tree.

To identify the *Cryptosporidium* species present in the samples, we performed nested PCR within the 18S rRNA gene on the 9 samples and Sanger sequenced the amplicons. However, only NMIMR11, UKP196, UKH101, EC1, EC4 and UKUB17 had good sequences (QV20+ ≥20 for majority of bases sequenced, with average signal to noise ratios for each base called >100). Partial 18S sequencing from these samples were aligned against 18S rRNA sequences from different species of *Cryptosporidium* obtained from NCBI to construct a maximum-likelihood phylogenetic tree (Fig. 4B). The phylogenetic tree demonstrated that EC1 is closely related to but distinct from *C. meleagridis* (AF112574), EC4 is related to but distinct from *C. canis* (AF112576), NMIMR11 and UKP196 clade with *C. parvum* (AF108864), UKH101 to *C. hominis* (DQ286403.1) whereas UKUB17 is closely related to but distinct from *C. ubiquitum* (HM209366.1).

We next performed CES-Seq to assemble the genomes of the 9 *Cryptosporidium*-positive human clinical samples collected from different regions of the world (Fig. 4A). After capture hybridization, we produced libraries for each sample that were sequenced using the Illumina platform. All paired-end reads were filtered and aligned against the *C. parvum* Iowa-ATCC v54 reference genome to determine read coverage and read distribution. Two samples EC4 (*C. canis*) and FEgypt had only 2141 (0.014X coverage) and 103,590 (0.98X coverage) reads map to the reference genome, respectively (Table 3). Whereas samples C2 and C8 each had greater than 500,000 reads map (2.12X, 2.14X coverage, respectively), however, a majority of the reads from the sequencing run did not map, indicating a lower yield of *Cryptosporidium* DNA from the capture hybridization reaction (Table 3). In contrast, samples NMIMR11, UKUB17, UKH101, UKP196 and EC1 each possessed higher coverage, ranging from 4.7x to 531X (Table 3) with the number of reads mapping specifically to the *C. parvum* reference genome ranging from a low of 2.4% (UKUB17) to a high of 99% (NMIMR11). Importantly, for samples EC1, EC4, and FEgypt, which were coinfected with *Giardia* (C_T_ = 28.3, 28.6, 27.7, respectively), essentially no reads mapped to *Giardia,* confirming the specificity of the probes and the fidelity of the CES-Seq method to specifically pull out *Cryptosporidium* gDNA (Table 3). While the coverage and percentage of mapped reads generally tracked with the qPCR C_T_ (lower C_T_ corresponded to higher coverage and percentage of mapped to unmapped reads), this was not always the case, see for example sample EC4 that had a C_T_ of 29.47, but only 0.014X coverage versus sample FEgypt with a C_T_ of 35.46 but 0.979X coverage (Table 3); or sample EC1 with a C_T_ of 27.15 and read coverage of 85.56x versus sample C2 with a similar C_T_ of 26.46 but only 2.12X coverage. How sample storage/preservation (*i.e.,* freezing, ethanol fixation) or the species of *Cryptosporidium* present influences the success of the CES-Seq methodology is currently being evaluated, but may represent a significant variable that needs to be factored when performing this technique.

**Table 3.** Sample information.

| ID | Sample names | Country | Origin | Material | Number of reads | Mapped reads | Unmapped reads | Coverage | Ct for *Cryptosporidium* | 18S Type | Sequencer | Popscicle Type | Ct for *Giardia* | FastQ Screen Mapped to *G. lamblia* WB |
| --- | --- | --- | --- | --- | --- | --- | --- | --- | --- | --- | --- | --- | --- | --- |
| Dg045 | C2 | Ghana | Human | Stool | 88,344,601 | 622,429 / 0.7% | 87,722,172 / 99.3% | 2,1251 | 26.46 | *C. hominis* | MiSeq+NextSeq | *C. parvum* |  |  |
| Dg083 | C8 | Ghana | Human | Stool | 88,437,644 | 539,311 / 0.61% | 87,898,333 / 99.39% | 2,1369 | 28.87 | *C. hominis* | MiSeq+NextSeq | *C. parvum* |  |  |
| ENRH_0052 | NMIMR11 | Ghana | Human | Stool | 51,262,478 | 50,546,533 / 98.6% | 715,945 / 1.4% | 531,4183 | 17.19 | *C. parvum* | MiSeq | *C. parvum* |  |  |
| UKUB17 | UKUB17 | UK | Human | Stool | 27,999,948 | 668,726 / 2.39% | 27,331,222 / 97.61% | 4,7236 | 29.37 | *C. ubiquitum* | HiSeq | Different ancestral |  |  |
| UKH101 | UKH101 | UK | Human | Stool | 187,665,611 | 12,512,057 / 6.67% | 175,153,554 / 93.33% | 134,382 | 22.6 | *C. hominis* | MiSeq+NextSeq | *C. hominis* |  |  |
| UKP196 | UKP196 | UK | Human | Stool | 45,329,037 | 12,234,615 / 26.99% | 33,094,422 / 73.01% | 181,1639 | 25.85 | *C. parvum* | HiSeq | *C. parvum* |  |  |
| EC1 | EC1 | Ecuador | Human | Stool | 20,872,547 | 8,339,764 / 39.96% | 12,532,783 / 60.04% | 85,5575 | 27.15 | *C. meleagridis* | MiSeq | Different ancestral | 28.3 | 0 |
| EC4 | EC4 | Ecuador | Human | Stool | 136,152 | 2,141 / 1.57% | 134,011 / 98.43% | 0,0139 | 29.47 | *C. canis* | MiSeq | Not used | 28.6 | 6 |
| 51 Egypt OC | FEgypt | Egypt | Human | Stool | 3,196,662 | 103,590 / 3.24% | 3,093,072 / 96.76% | 0,9797 | 35.46 | none | MiSeq | Not used | 27.7 | 1 |

After reference mapping the reads, we used the Genome Analysis Toolkit (GATK) to call SNP variants across the genomes of the seven clinical samples that had coverage equal to or greater than 2X. SNP variants were filtered to select for only high confidence SNPs defined as those SNPs that were supported by at least 5 reads in at least one of the 7 samples in the VCF (see Methods). We next constructed hierarchy-based phylogenetic trees using only the high confidence SNPs identified after reference mapping the CES-Seq biological samples above against SNPs identified in the reference sequences that were comprised of 12 whole-genome sequences from different human-infective species that have been published previously (SRA database), specifically *C. parvum* (ERR1035621, ERR1738341 and ERR1760143), *C. hominis* (ERR311205, ERR311209 and ERR363534), *C. meleagridis* (SRR1179185 and SRR793561), *C. ubiquitum* (SRR7895345 and SRR7895268), *C. cuniculus* (SRR6813608) and *C. tyzzeri* (SRR5683558). The raw reads from these reference samples were likewise mapped against the *C. parvum* Iowa-ATCC v54 genome to call SNP variants. However, due to a wide range in the distribution and read coverage between the CES-Seq samples and reference samples, too few high confidence SNPs were identified when samples were analyzed altogether. Hence, it was more informative to produce individual trees for each CES-Seq sample. To do this, we developed individual SNP files after reference mapping each CES-Seq sample to increase the total number of high confidence SNPs (without any gaps) available for phylogenetic classification of each biological sample against a reference set of previously published genomes (Table S4). For EC1 329,989 high quality SNPs were identified to construct a phylogenetic tree using the reference genomes. Correspondingly, 432,372 SNPs were identified for UKP196, 58114 SNPs for UKUB17, 329989 SNPs for EC1, 24,716 SNPs for C2, and 14,582 SNPs for C8 network trees (Fig. 4C, Suppl. Fig. 2). This approach better resolved the phylogenetic position of each captured genome dataset within the context of different *Cryptosporidium* species to visually depict their evolutionary history and determine the extent to which recombination played a role in their origin. Notably, the hierarchy trees were largely congruent with the 18S rRNA maximum-likelihood tree in Fig. 4B, specifically, that UKP196 and NMIMR11 clustered closely with *C. parvum*, UKH101 with *C. hominis*, whereas EC1 was distinct from *C. meleagridis*, and UKUB17 was distinct from *C. ubiquitum* at WGS resolution and was recombinant (Fig. 4C). Of note, the *C. hominis* sample ERR363534, with Biosample ID ERS226604, clustered with *C. parvum* (Fig. 4C, Suppl. Fig. 2). The most parsimonious explanation for this result is that it was named erroneously in the Wellcome Pilot Study performed to sequence diverse *Giardia* and *Cryptosporidium* isolates.

### Genetic Population Structure of Human Clinical Samples

To produce a robust model for the evolutionary history and population genetic structure fingerprint for each clinical sample that underwent CES-Seq, it is imperative to know the relative read distribution, coverage, and whether any bias was introduced during the amplification step after capture hybridization. This is to ensure that all SNP variants called are of high confidence to support the genetic models and to inform on the possibility to perform GWAS studies using sequence datasets from this technique. To visualize this, we developed a number of custom analysis pipelines to generate SNP density statistic plots for the datasets assembled. For each of the CES-Seq datasets from the clinical samples investigated, we generated read depth statistics using SAMtools and plotted these values in in order to visualize 1) the median SNP density per chromosome, 2) the read depth per 10kb window, 3) the read coverage percentage per 10kb window, and 4) the normalized SNP density per 10kb window (Fig. 5). To confirm the utility of these pipeline scripts to visualize SNP distribution and depth across all 8 chromosomes in 10kb windows, we utilized the artificial *C. parvum*_1ng Bunchgrass sample mixture in human stool gDNA as a control for the analysis pipeline. All CES-Seq Illumina reads were first mapped to the *C. parvum* Iowa-ATCC v54 genome. Read depth per 10kb was at or above 50X and this was depicted using a green line (Fig. 5) except in a very few focal regions on Chromosomes 2, 3, 4, 5, 6, and 8 (comprised of less than 100kb of sequence total) where read depth dropped to 10-30X, which corresponded to regions mis-assembled between the Iowa II v34 (the genome used to pick RNA baits) vs. Iowa-ATCC v54 (the genome used for reference mapping CES-Seq datasets), or undergoing copy number variation (Fig. 2D, 2E). Differences in the Bunchgrass vs. Iowa-ATCC genome may also result in a failure of some CES-Seq probes to capture sequences within these variant regions. Identification of genome blocks that contain high confidence SNPs is tantamount to facilitate prospective GWAS studies. To identify the genome-wide distribution and percentage of SNPs that are high confidence for each CES-Seq dataset, we calculated read coverage (depicted with a grey line) per 10kb to identify the percentage and distribution of SNPs that are considered high confidence based on read depth within each 10kb block (Fig. 5). Then, for each 10kb window, a normalized SNP density was calculated and plotted to identify the number of SNPs in each block that are both high confidence, and different from the reference genome, which were plotted using a blue dot. Finally, the red line for each chromosome represents the median SNP density compared to reference for each CES-Seq dataset, to estimate the extent of diversity from the reference genome, per chromosome. The integration of these SNP density statistic plots generates a confidence statistic for each dataset to inform on the degree of admixture, potential for mixed infection, and divergence from reference genomes for each CES-Seq dataset of *Cryptosporidium.* This methodology was then applied to each CES-Seq dataset for each of the 9 clinical samples investigated. *Cryptosporidium parvum* sample NMIMR11 from Ghana was highly similar to the Bunchgrass isolate, but clear differences in the SNP statistic per 10kb (blue dots) were readily visualized (Fig. 5). Additional CES-Seq datasets, including UKP196, UKH101, UKUB17, and EC1 each possessed sufficient read depth and read coverage (%) to identify high-confidence SNPs that were distributed genome-wide. Importantly for each 10kb window, there were sufficient variant SNPs detected that were distributed genome-wide to confer confidence that phylogenomic analyses to determine the evolutionary history, degree of admixture, and the ability to distinguish single strain from mixed infections is possible using the CES-Seq method. Further, for samples where genome-wide coverage was low - between 0.5-5X (samples FEgypt, C2, C8) - high confidence SNPs were still identified genome-wide, with sufficient density to infer phylogenetic ancestry. Only the EC4 CES-Seq dataset was insufficiently resolved to facilitate onward phylogenomic characterization (Fig. 5).

**Fig. 5.**
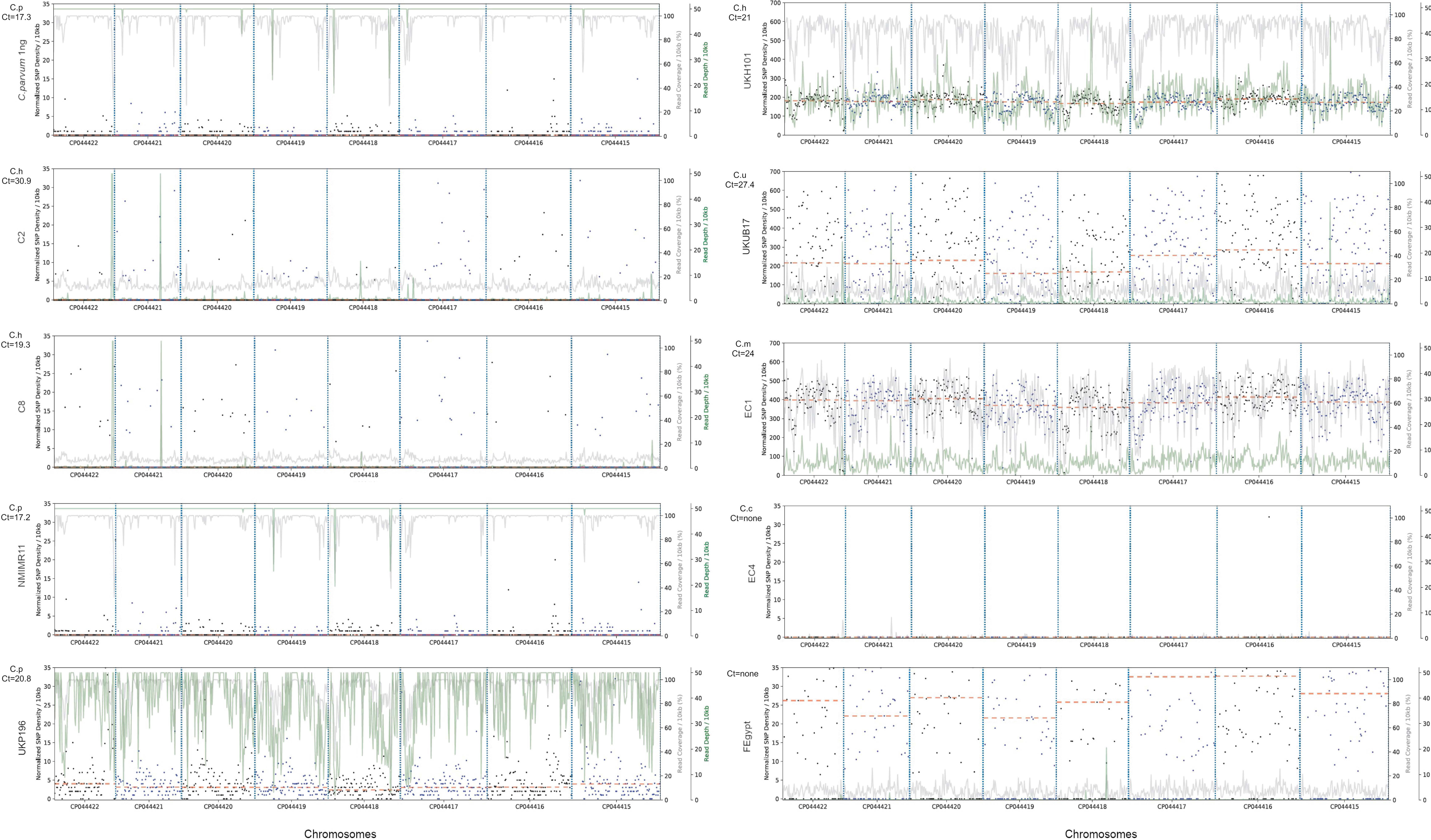
SNPs distribution of *Cryptosporidium* species directly from patient’s stool samples (C2, C8, NMIMR11, UKP196, UKH101, UKUB17, EC1, EC4 and FEgypt) and the *Cryptosporidium parvum* spiked control (*C. parvum* 1ng). SNP density plots were generated using an in-house python script. Blue dots represent normalized SNP density values by 10kb. Green line, read depth / 10kb; gray line, read coverage / 10kb; red dashed line, median SNP density per chromosome (not considering sequencing gaps).

To determine the ancestry and relatedness within the population genetic structure of *Cryptosporidium* for each CES-Seq genome level dataset, we first needed to resolve whether the high confidence SNPs identified predicted a mixed-species infection. We used DEploid to identify the number of haplotypes and their relative frequency present in each CES-Seq dataset. Each VCF file was mapped against a reference PANEL of 12 *C. parvum*, 8 *C. hominis*, 2 *C. meleagridis,* 2 *C. ubiquitum*, and 1 *C. tyzzeri* genomes (Table S4) to determine weight supported allele frequencies, plotted as histograms, to resolve the number of haplotypes present, their relative frequency, and hence, whether each CES-Seq dataset was from a single isolate. As proof-of-principle, we re-ran the DEploid analysis pipeline using the raw read sequences from the mixed species mock community generated by spiking an equivalent amount of *C. parvum, C. hominis,* and *C. meleagridis* into human fecal DNA, capture hybridized, sequenced and mapped against the updated reference genome CryptoDB-54_CparvumIowa-ATCC_Genome. As in Figure 3B, 3 haplotypes were resolved with a ratio of 58% *C. parvum*, 32% *C. hominis,* and 12% *C. meleagridis* (Fig. 6A, top left panel). We then assessed each CES-Seq dataset for NMIMR11, UKP196, UKH101, UKUB17 and EC1, samples that possessed sufficient read coverage and depth to call high confidence SNPs. DEploid predicted that all 5 were single isolate infections (Fig. 6A). Although NMIMR11 did possess a low frequency of minor alleles, they were too few, and likely represent genetic drift or could possibly represent a mixed strain infection between two highly similar *C. parvum* isolates that are not resolved by this analysis, because the vast majority of SNPs mapped to a single *C. parvum* haplotype (Fig. 6A).

**Fig. 6.**
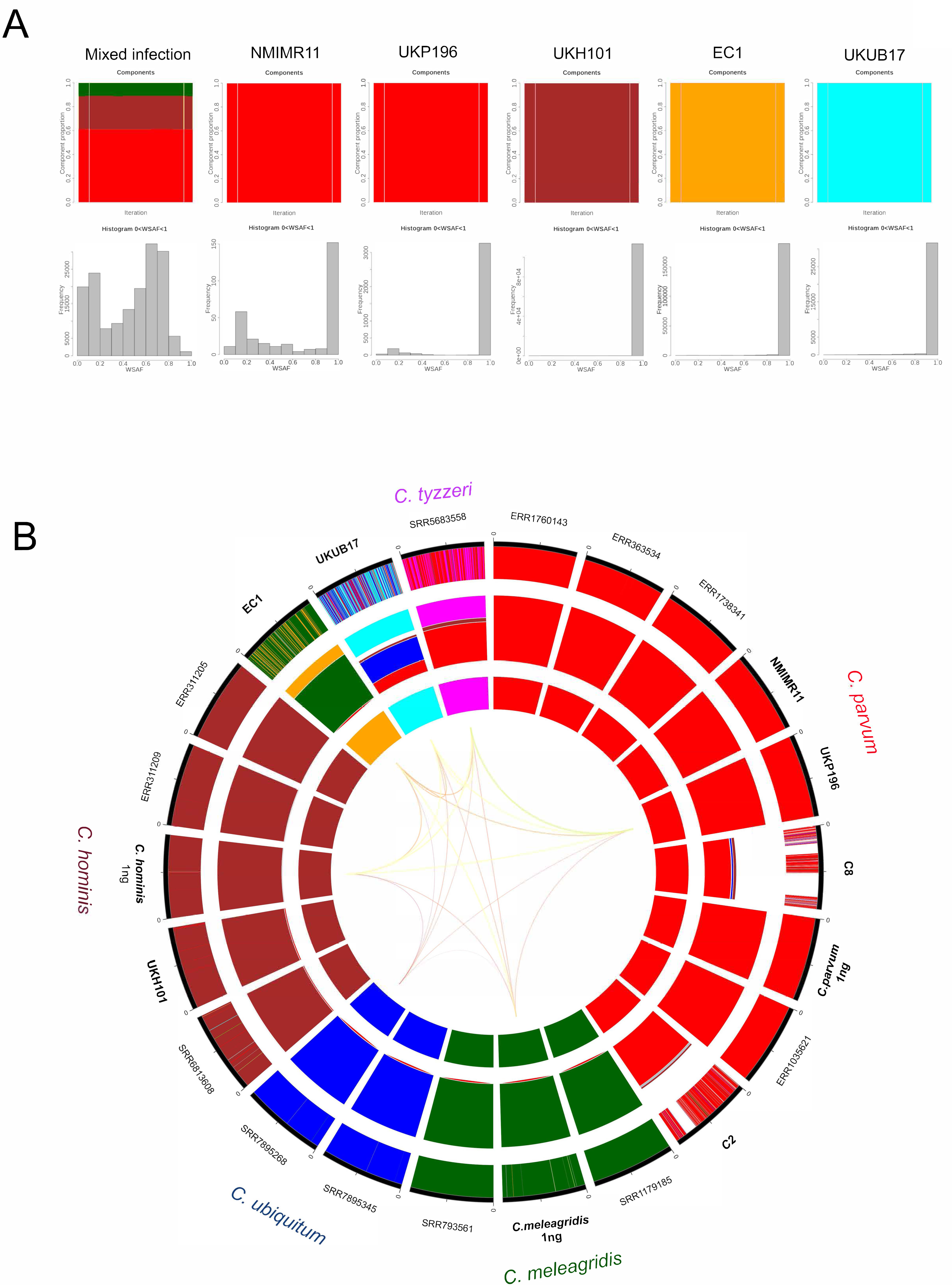
SureSelect-WGS population genetic structure of *Cryptosporidium* species. A) DEploid software was used to determine if the clinical sample was from a single strain, or mixed strain infection. Genome-level datasets were mapped against a PANEL of 25 different reference *Cryptosporidium* species (Table S4) to determine the relative proportion of different haplotypes present in mixed-strain clinical samples and plotted as bar plots. The relative proportion of each genotype was pseudocolored to reflect the underlying genome present. The following pseudocolor code based on Fig. 6B was used: red for *C. parvum*; green for *C. meleagridis*; brown for *C. hominis*; blue for *C. ubiquitum*; orange for EC1-like, and cyan for UKUB17-like. The histograms in gray represent the weight supported allele frequencies (WSAF) for each clinical sample. Single histograms with WSAF values between 0.9-1.0 indicate a single strain infection, and the bar plot is pseudocolored to reflect the species genotype resolved. B) Circos plot representation of admixture analysis and chromosome painting of *Cryptosporidium* conducted by POPSICLE (6) based on an ancestral population size of K=7. Outermost concentric circle represents a chromosome-level admixture profile for each strain, painted in 10 kb sliding windows and pseudocolored to represent the species type resolved. The middle concentric circle plots the percentage of shared ancestry. The innermost concentric circle indicates the cluster assignment by color hue for a population size of K=7. Each color represents one ancestral population. Representative sequences were downloaded from SRA (https://www.ncbi.nlm.nih.gov/sra). Whole genome sequences of the clinical samples (yellow dots) were obtained after CES-Seq.

We next estimated the number of supported ancestries (K) that could be resolved among the five high confidence CES-Seq genome-level datasets for NMIMR11, UKP196, UKH101, UKUB17 and EC1, as well as the lower coverage datasets for C2 and C8 compared against representative genomes of 4 *C. parvum*, 3 *C. hominis*, 2 *C. meleagridis, 2 C. ubiquitum,* and 1 *C. tyzzeri* downloaded from the SRA. As controls, we also included the CES-Seq datasets generated using 1ng of *C. hominis* (TU502 isolate), 1ng of *C. parvum* (Bunchgrass isolate) and 1ng of *C. meleagridis* (TU1867 isolate). To do this, we calculated the Dunn index (61), which supported K=7 ancestral populations for the 22 genomes analyzed against each other. To visualize the shared ancestry across the different isolates and species of *Cryptosporidium* analyzed, we used the POPSICLE software to cluster the genomes into seven different colors and distribute them within the inner ring of the circos plot (Fig. 6B). POPSICLE also calculated the number of clades present for every 10 kb window across each input genome, and each clade was assigned a different color hue that was painted across the genome to resolve ancestry and the degree to which recombination had impacted each isolate. This was displayed in the outer ring of the circos plot (Fig. 6B). The middle ring of the circos plot estimated the percentage of each ancestry, represented by a different color hue, present in each genome-level dataset. Among the genomes downloaded from the SRA database, POPSICLE assigned the predicted species type recorded in the SRA, except for ERR363534 that was listed as *C. hominis*, but our POPSICLE and network tree analysis rather showed that it shared ancestry with *C. parvum*. Also, the highly similar *C. cuniculus* genome (SRR6813608) clustered together with *C. hominis* at K=7 and was only resolved from *C. hominis* at K=8. As expected, all three CES-Seq datasets from the artificial mixture samples successfully clustered together with their respective species types (*C. parvum, C. hominis or C. meleagridis*). Importantly, the C2 and C8 partial genomes resolved unambiguously with *C. parvum*. These specimens failed to genotype using sequencing primers targeting the sensitive 18S rRNA locus, but the CES-Seq partial genome datasets facilitated species assignment, highlighting the sensitivity of the CES-Seq method.

POPSICLE identified mosaic ancestries for two human isolates EC1 and UKUB17, that had been previously genotyped at the 18S rRNA locus as *C. meleagridis* and *C. ubiquitum*, respectively. However, genome-wide, EC1 resolved as a genetic mosaic, that shared 80.75% of its genome with *C. meleagridis* (darkgreen) and 18.70% of its genome with a new ancestry (orange) that had introgressed throughout the genome in large haplotype blocks that resembled genetic recombination (middle ring, circos plot). Likewise, UKUB17 shared 37.1% of its genome with *C. ubiquitum* (blue), however, 18.5% of the genome resolved with *C. parvum* (red), 3.5% with *C. hominis* (brown), and 34.2% with a distinct, new ancestry (cyan). In addition, 6.7% of the assembled genome had no reads that mapped to the *C. parvum* reference (white). These regions were distributed in discrete haploblocks throughout the genome and were largely restricted to sub-telomeric and telomeric sites (Fig. 6B). DEploid analysis unambiguously assigned these two assemblies as single isolates, suggesting that the two isolates recovered from two individuals likely represent inter-specific recombinants and highlight the utility of the CES-Seq method to inform on the genetic diversity within zoonotic species of *Cryptosporidium*.

To further resolve the ancestry and distribution of the predicted admixture blocks identified in the POPSICLE analysis for EC1 and UKUB17, we developed a new software pipeline that calculated the percentage of different SNPs within non-overlapping sliding windows of 10kb between each CES-Seq dataset compared against closely related species of *Cryptosporidium.* These values were then plotted across each chromosome to resolve ancestry and transitions consistent with recombination breakpoints (Fig. 7 and Suppl. Fig. 3 to 5). For EC1 and NMIMR11 (control) we used C*. parvum, C. meleagridis* and *C. hominis* genome references; for UKUB17 we used C*. parvum, C. hominis and C. ubiquitum* genome references because this sample was predicted to be *C. ubiquitum* by typing at the 18S rRNA locus and possessed a proportion of *C. ubiquitum* ancestry by POPSICLE (Fig. 6B). Pair-wise SNP plots for NMIMR11 resolved this sample as *C. parvum* genome-wide (magenta line for *C. parvum* mapped at essentially zero percent of different SNPs per 10 kb window) with no evidence of recombination. In contrast, EC1 was polymorphic, possessed clear ancestral blocks related to *C. meleagridis* (blue line mapped close to zero percent of different SNPs per 10kb window), however it also possessed large haploblocks that were divergent and unrelated to *C. parvum* (magenta line) or *C. hominis* (green line). In the genome-wide plots, there was also evidence of 7 haploblocks that were in fact *C. hominis* or *C. parvum*, suggesting that inter-specific recombination had impacted the population genetics of this human infective isolate. The pair-wise SNP haplotype plots also established that UKUB17 is novel, it was neither C*. parvum, C. hominis* nor *C. ubiquitum*, it was extensively recombined, with distinct haploblocks indistinguishable from *C. parvum, C. hominis,* or *C. ubiquitum* introgressed throughout the genome. It also possessed large haploblocks that were new, that did not BLAST with any known species of *Cryptosporidium* thus far analyzed. The blocks were as similar to *C. ubiquitum* as *C. hominis* is to *C. parvum,* which mapped to NMIMR11 genome-wide in the 10-20 percent of different SNPs per 10 kb, which may suggest that the differences are significant enough to reflect introgression of a new species-type thus far not resolved in the population genetics of *Cryptosporidium* (Fig. 7 and Suppl. Fig. 3 to 5). The DEploid analysis unequivocally predicted UKUB17 was the result of a single clone infection. The pair-wise SNP haplotype plots likewise showed that it was not synonymous with a mixed infection, as it produced unambiguous recombination break-points. Rather, the data and analysis pipelines herein supported UKUB17 to be an inter-specific genetic mosaic of mixed ancestry within a single haplotype.

**Fig. 7.**
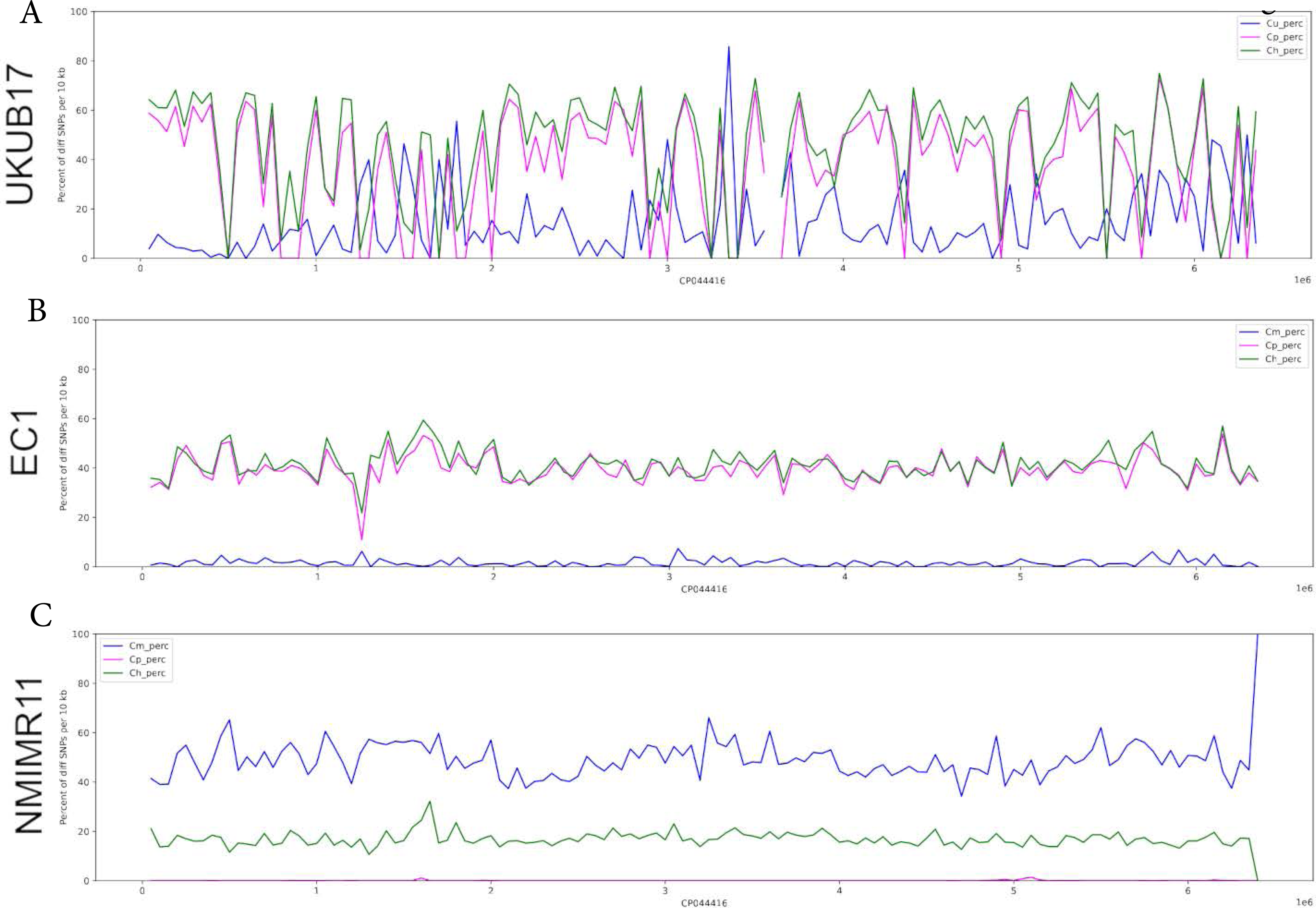
Pair-wise SNP haplotype plots used to resolve the degree of identity of each CES-Seq examined clinical sample against reference genomes from related *Cryptosporidium* species. The percentage of SNPs different between the strain of interest (*i.e.,* Isolate UKUB17, EC1, or NMIMR11) and the reference strains of either *C. parvum* (magenta line), *C. meleagridis* (blue line), *C. hominis* (green line) or *C. ubiquitum* (blue line) were calculated and plotted in non-overlapping sliding windows of 10kb along each chromosome (x-axis, chromosome position in Mb). Colored lines at the baseline of “0” percent of different SNPs per 10kb indicated that the clinical sample had 100% identity to the reference genome of that color type. A) UKUB17 sample, blue - *C. ubiquitum*, magenta – *C. parvum* and green – *C. hominis.* B) EC1 sample, blue - *C. meleagridis*, magenta – *C. parvum* and green – *C. hominis*.

## Discussion

In this study, we applied Capture Enrichment Sequencing (CES-Seq) to generate genome-level datasets of *Cryptosporidium* associated predominantly with asymptomatic carriage directly from human stool specimens. The majority of samples had been previously frozen or preserved in ethanol. We demonstrated that ∼75,000 RNA baits that cover ∼92% of the *Cryptosporidium* genome were sufficient to capture and enrich *Cryptosporidium* DNA ∼1500 fold directly from stool samples with C_T_ below 28, and for samples with C_T_ between 28 and 31, the ability to produce partial genomes for phylogenetic analysis and genotyping far exceeded current multi-locus typing methods for the genus. We showed that the CES-Seq genome-wide sequencing method is both highly sensitive and specific and can successfully resolve species of *Cryptosporidium* at whole genome resolution from parasite gDNA present in as little as 0.005% in human stool DNA. Further, the baits generated against *C. parvum* captured and produced genome-level datasets for more distantly related species without the requirement or need to purify oocysts, including *C. meleagridis, C. canis, C. ubiquitum* as well as interspecific recombinants present in clinical/field samples. We also showed the applicability of applying various phylogenomic pipelines, including DEploid, SplitsTree, and POPSICLE software suites to infer genetic diversity, degree of mixed infection, and the population genetic structure of *Cryptosporidium* species infecting predominantly asymptomatic patients. Our approach demonstrates the utility and cost-effectiveness of *“in situ*” CES-Seq to generate genome level datasets of *Cryptosporidium* from patient samples to effectively genotype the *Cryptosporidium* present in endemic settings at whole genome resolution, and how to use phylogenomic analyses to infer the potential for their zoonotic transmission, drug resistance, and capacity to cause new disease or alter host range. Ultimately this technology, which enables high-throughput sequencing of samples without the requirement to isolate oocysts, should empower scientists in lower-to-middle income countries (LMICs) where transmission is endemic to better understand the burden of *Cryptosporidium* in their countries and track how infection changes over time.

With approximately 22 species identified to infect people within the genus *Cryptosporidium*, many of which are zoonotic and possess broad host ranges based on recent molecular epidemiological studies (8), it is essential to determine the applicability of utilizing RNA baits designed against one reference genome (in this case *C. parvum*) to effectively capture and enrich for other species of *Cryptosporidium* that infect humans. Data herein established that these baits, which represent the first set of baits developed for the *Cryptosporidium* field, possessed a broad specificity, and were capable of pulling out, with high efficiency and at whole genome resolution, zoonotic species such as *C. meleagridis, C. canis*, and *C. ubiquitum,* as well as interspecific recombinants. However, the RNA baits in this paper were not designed to capture gene-poor sub-telomeric and telomeric regions, because they were designed prior to the release of the most up-to-date telomere-to-telomere (T2T) assembly that identified additional genes not present in the Iowa II v34 *C. parvum* assembly (65). Importantly, the CES-Seq datasets generated using baits designed against the first Iowa II v34 assembly did map with high fidelity to the new reference *C. parvum* T2T Iowa-ATCC v54 *C. parvum* assembly and our ability to perform robust population genomic analyses was not impacted. With the current push within the *Cryptosporidium* community to generate *de novo* genomes for a majority of the zoonotic species, future studies to increase the sensitivity of this approach should focus on generating additional RNA baits that span genomic regions that are either highly divergent in the other 21 species that infect humans or not present in *C. parvum*. One such effort that shows great promise recently used ∼ 10 genomes from different *Cryptosporidium* species to design a new bait set to more broadly detect diverse *Cryptosporidium* species and genotypes both *in silico* and experimentally for specificity and sensitivity (66).

Recent molecular epidemiological and genomic surveillance studies have established that the genus *Cryptosporidium* exhibits a complex evolutionary and transmission dynamics, including recombination events that contribute to adaptive genetic exchange between different species (interspecific) and within the same species (intraspecific) of *Cryptosporidium.* Such recombination events observed among zoonotic species of *Cryptosporidium* have highlighted the role of genetic hybridization in the emergence and enhanced transmission of specific genotypes that possess an increased capacity to cause disease or alter their host range (29–33). CES-Seq genome-level datasets sequenced directly from patient samples should greatly facilitate studies aimed at understanding the true genetic diversity of *Cryptosporidium* species and the role of recombination in the emergence of new genotypes associated with outbreaks, or that are circulating in endemic regions. Indeed, our NGS workflow identified extant recombination within the single UKUB17 specimen that typed as *C. ubiquitum* at the 18S rRNA gene (Fig. 4) but was shown to be a complex mosaic possessing large haploblocks with ancestry belonging to *C. parvum, C. hominis, C. ubiquitum,* and a novel sequence type (Fig. 7). This capacity to resolve genomes at high-resolution directly from human stool samples should inform future work that tests the hypothesis that the evolution of anthroponotic transmission in humans is the result of acquiring pathogenicity determinants from within the pan-*Cryptosporidium* genome to make human infection possible (30,31,33), analogous to the inheritance of secreted pathogenicity islands that determine host adaptation and disease potential in the related apicomplexan parasite *Toxoplasma gondii* (67,68). Importantly, the CES-Seq method and the associated population genomic pipelines developed herein also makes it relatively straightforward to determine the degree to which co-circulating strains (mixed infections) occur among susceptible hosts, including humans. This is necessary information, for two reasons: 1) the degree to which mixed infection occurs provides an estimate for the role genetic exchange plays in the evolution of genotypes that possess new host preferences or the capacity for zoonotic versus anthroponotic transmission (21); and 2), the ability to resolve whether the infection is by a single genotype is relevant for genotype to phenotype studies to support GWAS analyses. Hence, estimation of the rate and relatedness of such mixed infections is a critical factor to correctly construct the population genetic structure of *Cryptosporidium,* particularly in endemic regions, where the probability of mixed infection is relatively high (69,70).

In our study, the CES-Seq methodology coupled with the computational pipelines we applied and/or developed to analyze the assemblies produced establishes a new paradigm for resolving the genome sequences present in both archived and prospectively collected biological samples. Our workflow facilitates the identification of high confidence SNPs that are required for robust phylogenetic resolution of derived genome datasets in the context of defined *Cryptosporidium* species to determine the true population genetic structure of circulating strains, in reference to their zoonotic transmission. Our computational analysis platform is also capable of resolving mixed-strain infections, determining the genome structure of single strain infections, and applying new visualization tools to identify high-confidence SNPs in order to envisage future GWAS studies to map specific genes that contribute to phenotypic traits. Understanding the genetic basis of adaptation and admixture in *Cryptosporidium* populations is crucial for elucidating the epidemiology and pathogenesis of cryptosporidiosis, informing public health interventions, and guiding the development of effective control strategies.

In summary, the CES-Seq genome-level sequencing method is a novel, highly sensitive and specific tool to conduct phylogenomic studies on *Cryptosporidium* species circulating in stool samples to understand the population genetic structure of *Cryptosporidium* pan-genomes throughout the world. These studies are necessary to understand not only the zoonotic potential of a strain, but also the evolution and emergence of novel subtypes that impact *Cryptosporidium* transmission, host adaptation, and disease potential both locally (in the context of an outbreak) and globally (in the context of a selective sweep).

## Acknowledgements

The authors are supported by the Intramural Research Program of the National Institute of Allergy and Infectious Diseases (NIAID) at the National Institutes of Health. We thank Genomic Technologies Section at NIAID particularly Dr. Timothy Myers, Dr. Qin Su and Francisco Otaizo-Carrasquero for conduction Illumina sequencing and helpful comments. We also thank Dr. Jessica Kissinger and Dr. Travis Glenn for their support and helpful discussions throughout the production phase of this project. The following reagents were obtained through the NIH Biodefense and Emerging Infections Research Resources Repository, NIAID, NIH: Genomic DNA from *Cryptosporidium hominis*, Isolate TU502, NR-2520; *Cryptosporidium parvum*, Isolate Iowa, NR-2519; *Cryptosporidium meleagridis*, Isolate TU1867, NR-2521.

## Author contributions

Study was designed by AK, EVCAF and MEG. Experiments were conducted by AK, EVCAF, and HV. Samples were provided by MCP, SB, IA, GR and RMC. Analysis was conducted by AK, HL, EVCAF, and HV, and the manuscript was written by AK, EVCAF and MEG.

---

**Supplemental Fig. 1.**
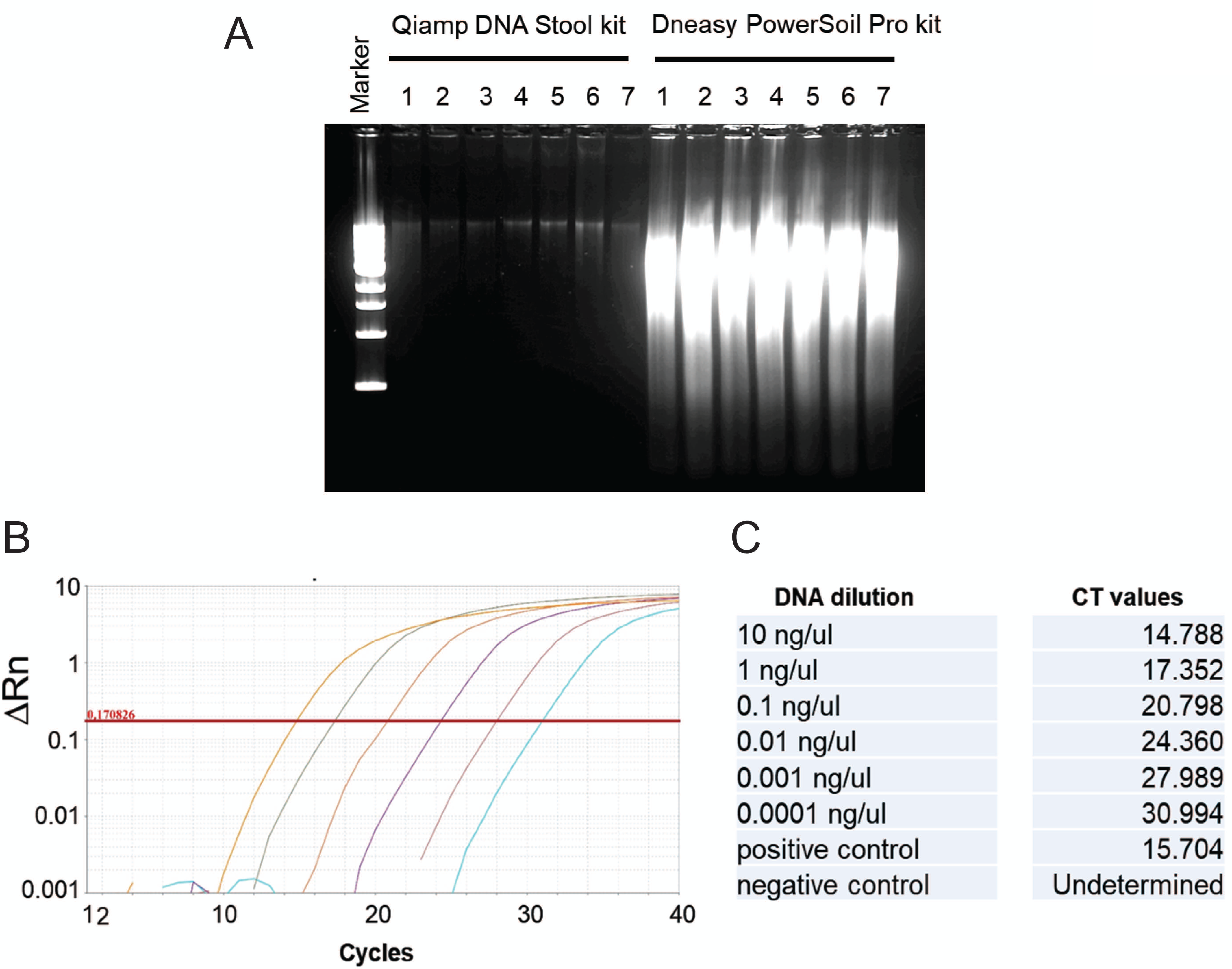
Sensitivity and Specificity Analysis of CES-Seq Method. A) gDNA extraction comparison using two different kits shows high yield using DNeasy PowerSoil Pro kit. B and C) 18S rRNA qPCR assay using Serial dilutions of Cryptosporidium parvum gDNA in human stool gDNA.

**Supplemental Fig. 2.**
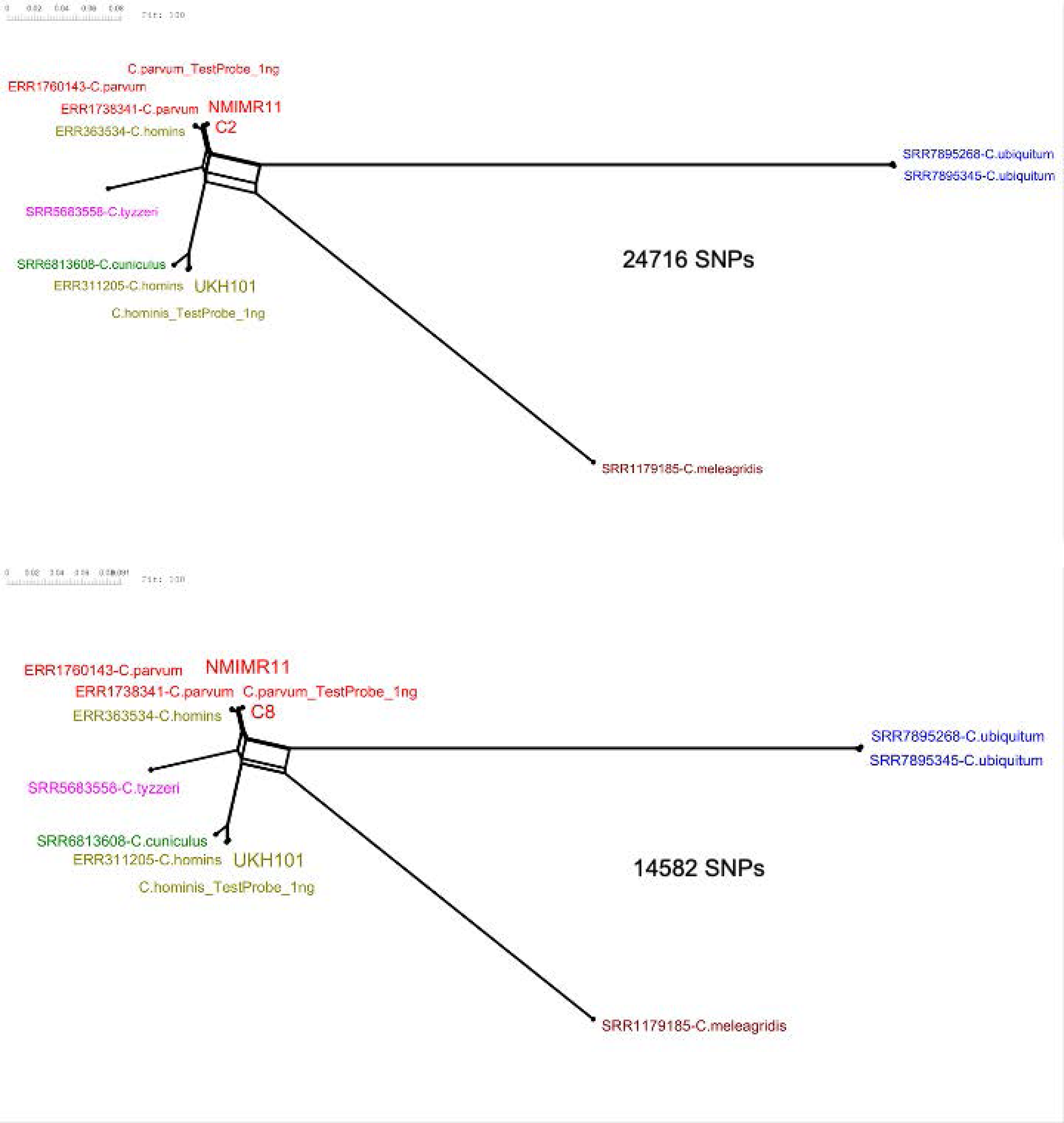
Phylogenetic Network Trees of C2 and C8 samples. Trees were generated using SplitsTree (v4.17.1), total number of SNPs are indicated below each tree.

**Supplemental Fig. 3.**
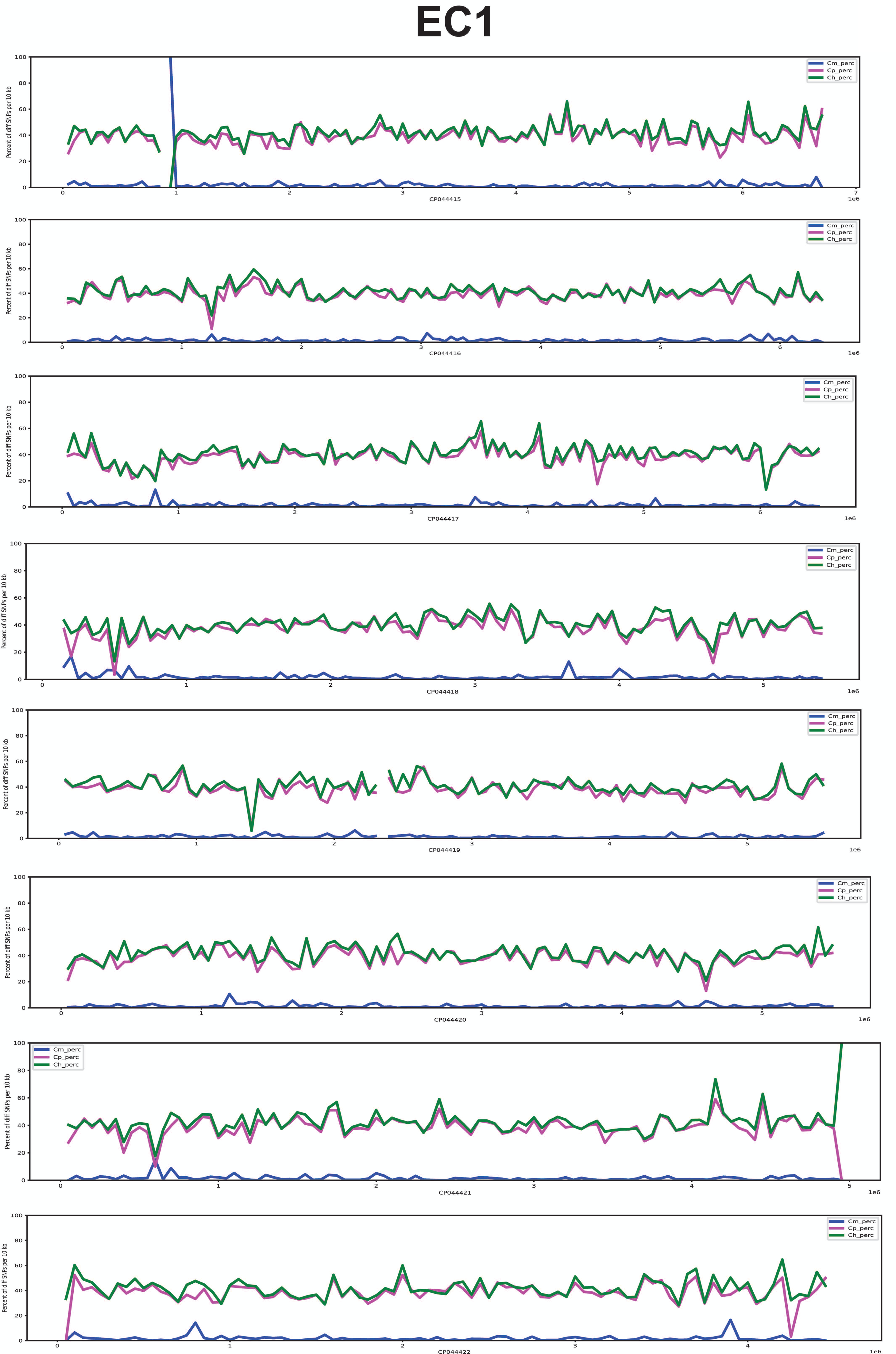
Pair-wise SNP haplotype plot (10kb sliding window) for all chromosomes for sample EC1.

**Supplemental Fig. 4.**
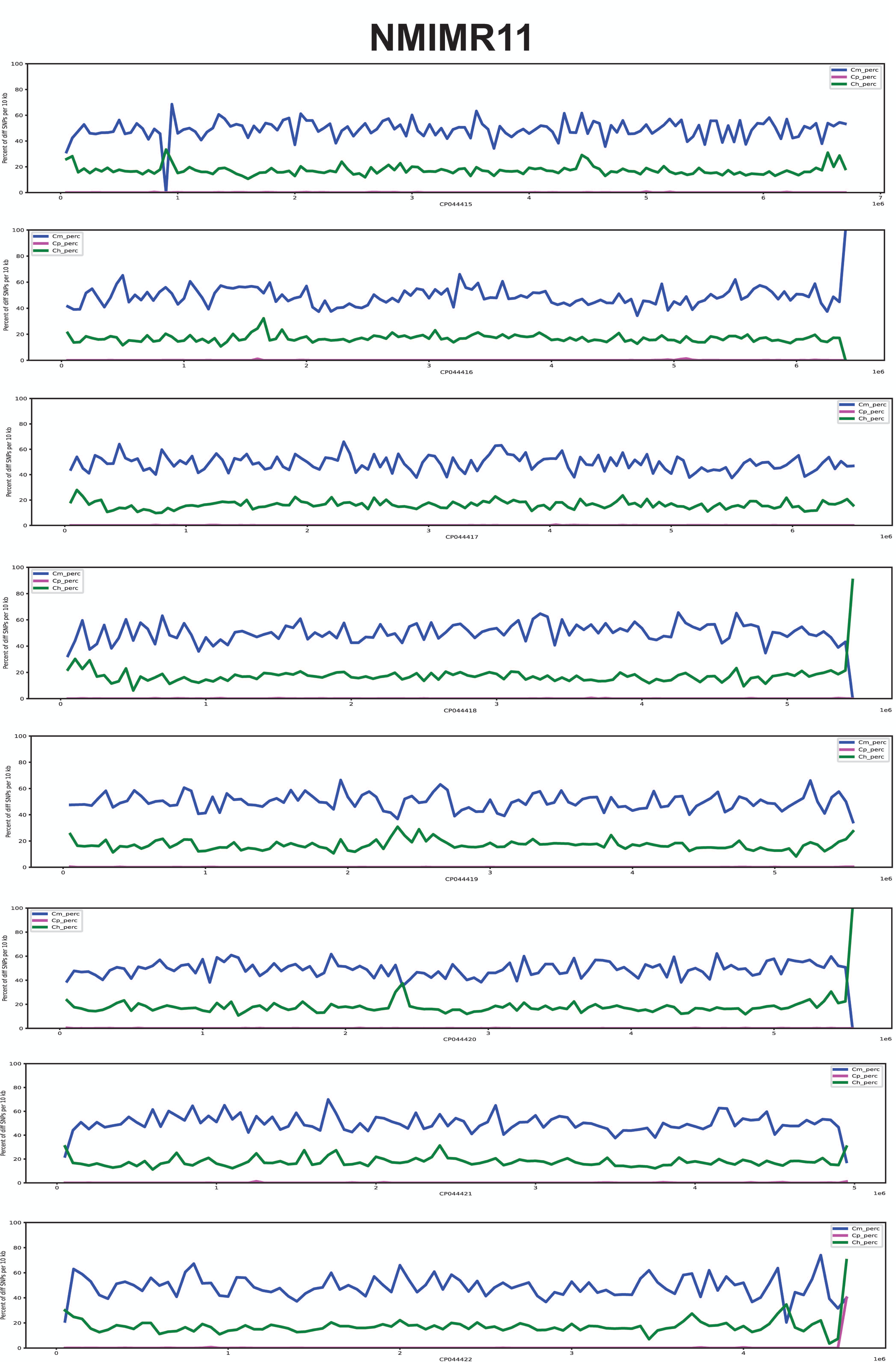
Pair-wise SNP haplotype plot (10kb sliding window) for all chromosomes for sample NMIMR11.

**Supplemental Fig. 5.**
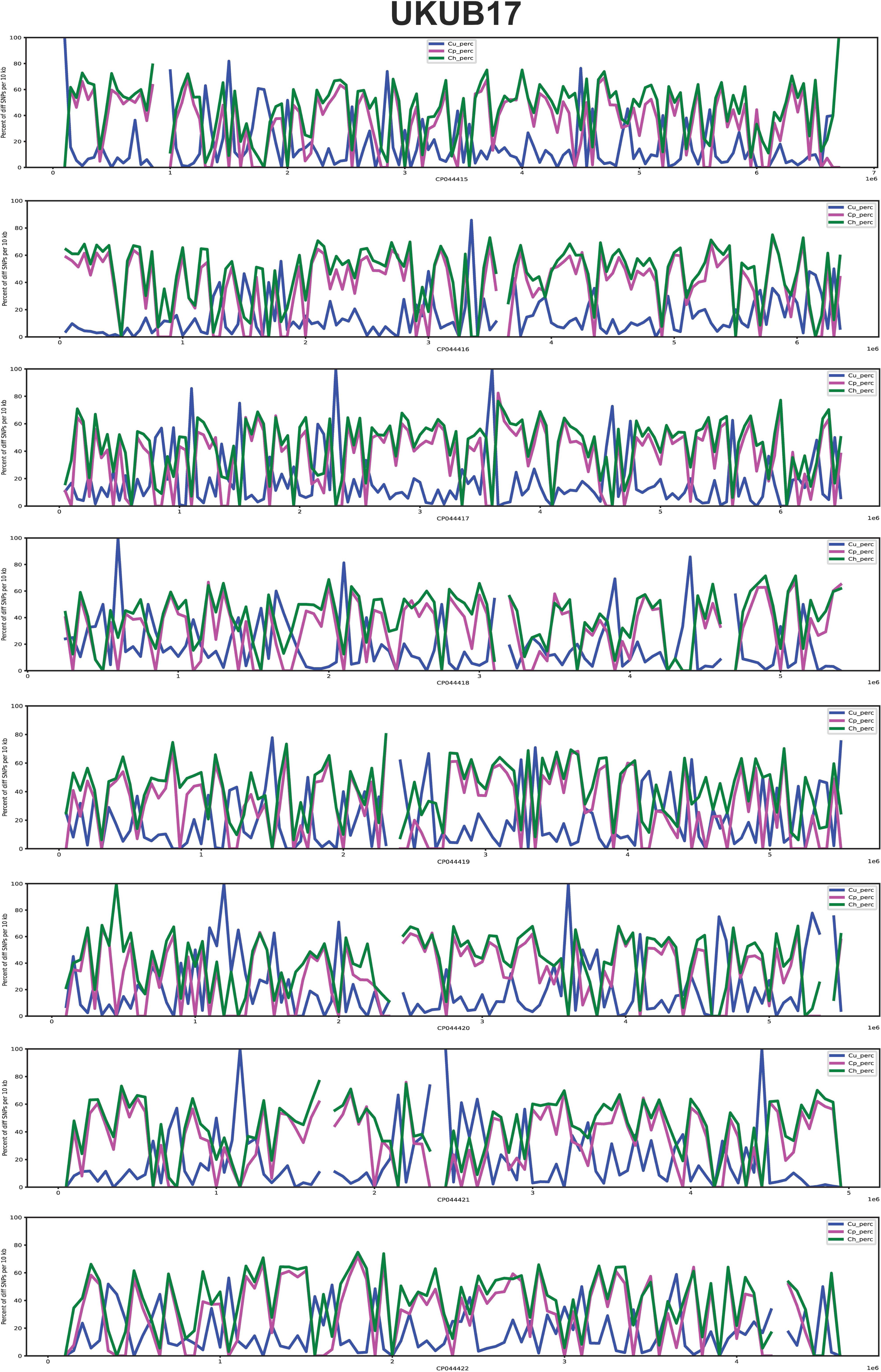
Pair-wise SNP haplotype plot (10kb sliding window) for all chromosomes for sample UKUB17.

**Table S1.**
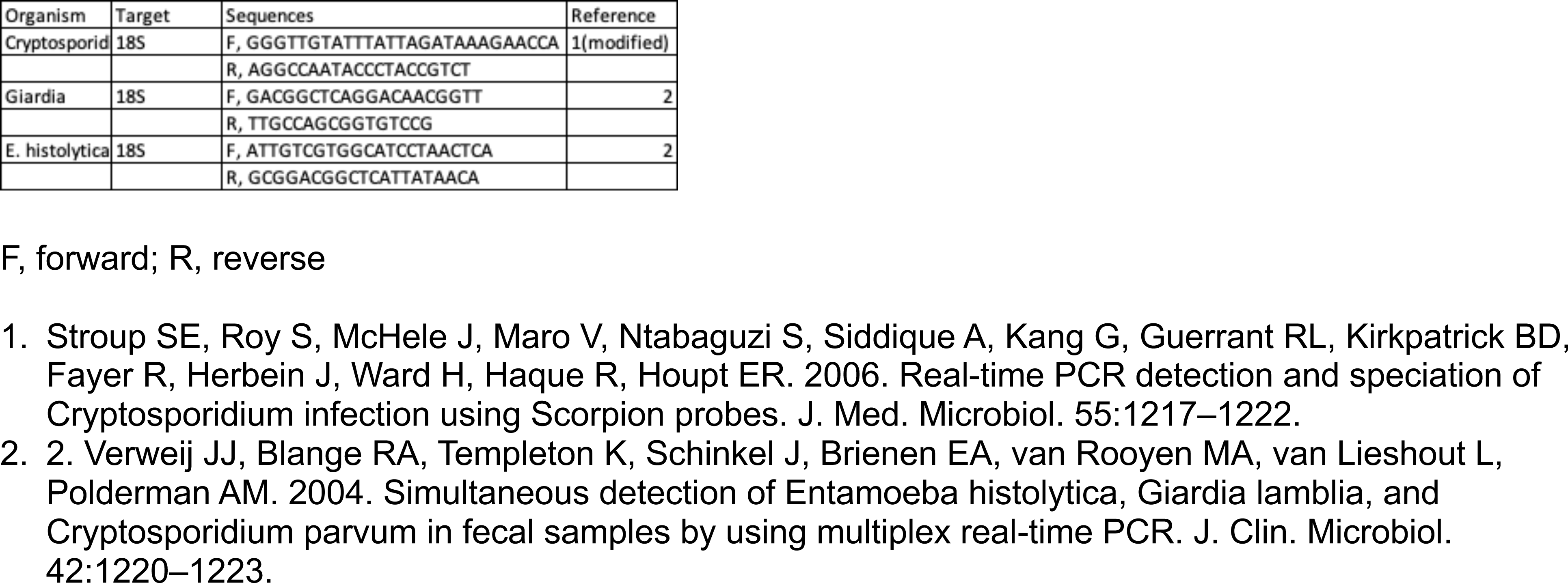
18S Ribosomal RNA primers used in this study.

**Table S2.**
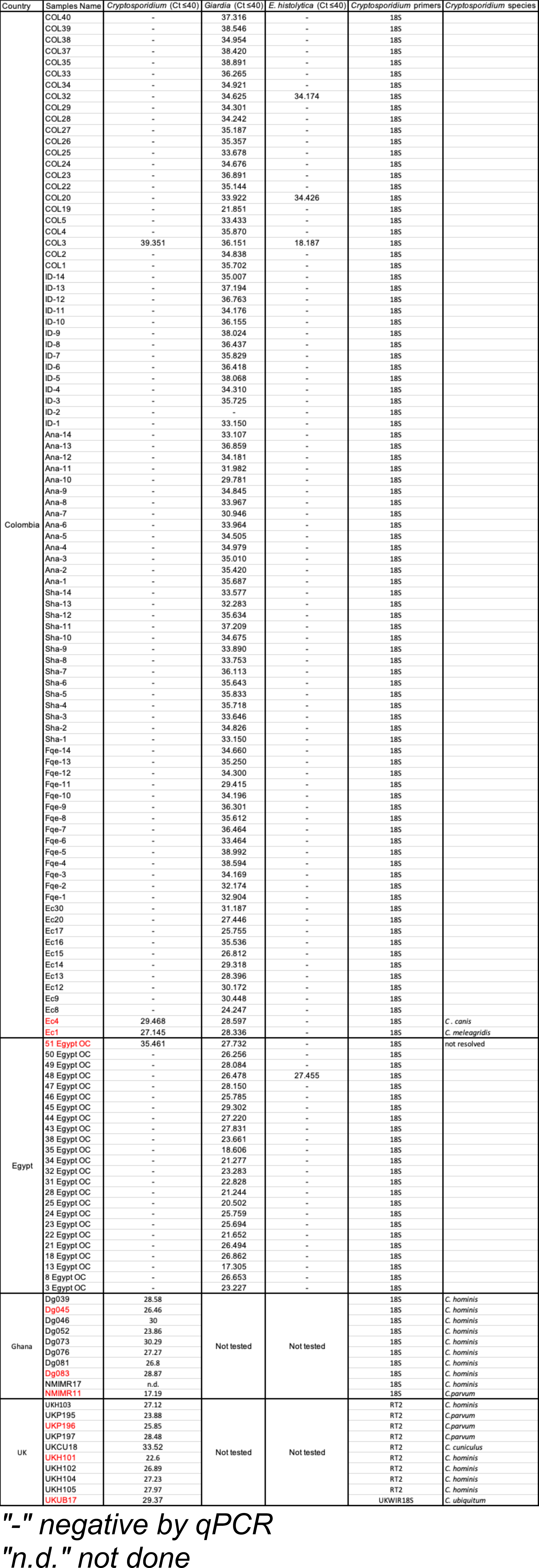
C_T_ values of 18S Ribosomal RNA qPCR assay using species specific primers for *Cryptosporidium*, *Giardia*, and *E. histolytica*.

**Table S3.** Chromosome ID for both genome references used.

| Reference used | Chromossomes ID | Chromossome Number | Chromossomes ID | Reference used |
| --- | --- | --- | --- | --- |
| *C. parvum* lowall<br>(CryptoDB Version 34) | CM000429 | 1 | CP044422.1 | *C. parvum* lowa-ATCC<br>(CryptoDB Version 54) |
|  | CM000430 | 2 | CP044421.1 |  |
|  | CM000431 | 3 | CP044420.1 |  |
|  | CM000432 | 4 | CP044419.1 |  |
|  | CM000433 | 5 | CP044418.1 |  |
|  | CM000434 | 6 | CP044417.1 |  |
|  | CM000435 | 7 | CP044416.1 |  |
|  | CM000436 | 8 | CP044415.1 |  |

**Table S4.** PANEL of 25 different reference *Cryptosporidium* species. Each VCF file was mapped against a reference PANEL of 12 *C. parvum* (ERR1035619, ERR1035621, ERR1738340, ERR1738345, ERR1738350, ERR1738337, ERR2366918, ERR961651, SRR6117460, SRR6147472, SRR6147945 and SRR614796), 8 *C. hominis* (ERR1305020, ERR1305025, ERR2240056, ERR2240065, ERR2240074, ERR2366927, ERR2366935 and SRR1015721), 2 *C. meleagridis (*SRR1179185 and SRR793561), 1 *C. tyzzeri (*SRR5683558) and 2 *C. ubiquitum* genomes (SRR7895268 and SRR7895345).

|  |  |
| --- | --- |
| ERR1035619 | *Cryptosporidium parvum* |
| ERR1035621 | *Cryptosporidium parvum* |
| ERR1738340 | *Cryptosporidium parvum* |
| ERR1738345 | *Cryptosporidium parvum* |
| ERR1738350 | *Cryptosporidium parvum* |
| ERR1738337 | *Cryptosporidium parvum* |
| ERR2366918 | *Cryptosporidium parvum* |
| ERR961651 | *Cryptosporidium parvum* |
| SRR6117460 | *Cryptosporidium parvum* |
| SRR6147472 | *Cryptosporidium parvum* |
| SRR6147945 | *Cryptosporidium parvum* |
| SRR6147964 | *Cryptosporidium parvum* |
| ERR1305020 | *Cryptosporidium hominis* |
| ERR1305025 | *Cryptosporidium hominis* |
| ERR2240056 | *Cryptosporidium hominis* |
| ERR2240065 | *Cryptosporidium hominis* |
| ERR2240074 | *Cryptosporidium hominis* |
| ERR2366927 | *Cryptosporidium hominis* |
| ERR2366935 | *Cryptosporidium hominis* |
| SRR1015721 | *Cryptosporidium hominis* |
| SRR1179185 | *Cryptosporidium meleagridis* |
| SRR793561 | *Cryptosporidium meleagridis* |
| SRR7895268 | *Cryptosporidium ubiquitum* |
| SRR7895345 | *Cryptosporidium ubiquitum* |
| SRR5683558 | *Cryptosporidium tyzzeri* |

